# PfK13-associated artemisinin resistance slows drug activation and enhances antioxidant defence, which can be overcome with sulforaphane

**DOI:** 10.1101/2025.10.05.680568

**Authors:** Ghizal Siddiqui, Carlo Giannangelo, Natalie A. Counihan, Yunyang Zhou, Annie Roys, Amanda De Paoli, Bethany M. Anderson, Matthew P. Challis, Lee M. Yeoh, Stuart A. Ralph, Laura Edgington-Mitchell, Tania F. de Koning-Ward, Darren J. Creek

## Abstract

Artemisinin resistance is globally prevalent, including in Africa, raising concerns and highlighting the need to better understand the cellular mechanisms behind this resistance. In *Plasmodium falciparum*, artemisinin resistance is primarily attributed to mutations in the PfKelch13 (PfK13) gene. In this study, we performed proteomic analysis on a range of sensitive and artemisinin-resistant parasites (both laboratory-generated and field isolates), revealing specific dysregulation of PfK13 protein abundance. Reduced PfK13 levels were linked to impaired hemoglobin digestion, decreased free heme levels, and consequently, decreased artemisinin activation. Artemisinin resistant parasites also exhibited elevated thiol levels, indicating a more reduced cellular state. Targeting the parasite redox capacity with sulforaphane potentiated artemisinin activity *in vitro* and in an *in vivo* rodent *Plasmodium berghei* model, offering a potential strategy to overcome resistance. Our findings provide critical insights into the molecular mechanisms of artemisinin resistance and suggest novel therapeutic interventions to restore drug sensitivity.

**One Sentence Summary: PfK13 mutations drive artemisinin resistance in *Plasmodium* parasites by enhancing antioxidant defences, which can be targeted by redox modulators such as sulforaphane.**

## Main text

Malaria is responsible for extensive mortality and morbidity in humans, with malaria incidence climbing from 221 million cases in 2019 to 263 million cases in 2023^1^. A major challenge in global malaria control is the emergence of resistance against most antimalarial drugs, including the artemisinins. Artemisinin-based combination therapy (ACT), combining a fast-acting artemisinin derivative with a longer-acting partner drug, is the current recommended first-line treatment for uncomplicated malaria caused by *Plasmodium falciparum*^2^. In recent years the emergence and spread of artemisinin resistance in South-East Asia, sub-Saharan Africa, India, South America and Papua New Guinea has contributed to treatment failures^1^.

Artemisinin’s mode of action involves a two-step mechanism, activation by heme released from parasite hemoglobin digestion, which catalyses the cleavage of artemisinin’s endoperoxide bond^3^, then the resulting carbon-centred radical kills the parasite by alkylating a large number of parasite molecules, with particular enrichment for proteins in redox pathways^4^. This alkylation of proteins involved in parasite redox homeostasis disrupts crucial parasite redox processes, contributing to the artemisinin mechanism of action^4^.

Artemisinin resistance is associated with non-synonymous mutations in the *P. falciparum* Kelch13 (PfK13) (PF3D7_1343700) gene^5^ encoding the PfK13 protein. Each of these mutations likely destabilise the PfK13 protein, as resistant clinical isolates and laboratory parasites carrying the resistance-associated PfK13 mutations R539T, C580Y, R561H and M579I contain reduced cellular levels of PfK13^6–12^. However, there has also been some inconsistency in the literature around whether different mutations in clinical isolates are associated with reduced amounts of PfK13 in ring-stage parasites^11^.

The molecular mechanism by which the PfK13 mutations confer artemisinin resistance in clinical isolates may involve multiple cellular and metabolic processes, including hemoglobin uptake and digestion, the unfolded protein stress response, mitochondrial functions, oxidative stress responses (including tRNA thiouridine modification reprogramming), and an enhanced DNA damage repair mechanism^3, 8–11, 13–20^. The PfK13 protein itself has been shown to be localised to cytostomes and involved in endocytosis of host cell cytosol in early ring stage parasites^8, 10, 15^. Endocytosis of red blood cell cytosol allows the parasite to catabolise host cell proteins into amino acids, a process critical for parasite *de novo* protein synthesis. As the main constituent of red blood cell cytosol is hemoglobin, artemisinin-resistant parasites take up less hemoglobin and have been shown to produce less redox-active ferrous heme, at least in mid-ring-stage parasites^21^. This was subsequently shown to be associated with lower levels of heme-artemisinin adducts^21^ and reduced fluorescence labelling of proteins using an artemisinin-based activity probe^12^. However, direct evidence for a decreased rate of artemisinin activation in artemisinin resistant early-ring stage parasites is yet to be shown.

In addition to acting as the major intracellular activator of artemisinins, ferrous heme is the primary source of oxidative stress during the blood stage of *P. falciparum*^22^. The parasite has established distinct mechanisms to detoxify heme including: (1) autooxidation to the non-toxic ferric state, producing reactive oxygen species (ROS)^22^, (2) polymerisation to non-toxic hemozoin^23, 24^, and (3) degradation by glutathione (GSH)^25^. We have shown previously that artemisinin-resistant parasites have increased levels of GSH, the parasite’s predominant reduced cellular thiol^9^. PfK13 mutations have also been shown to be associated with an increased baseline stress response^16, 17, 20^. Based on the impact of PfK13 mutations on hemoglobin uptake and GSH levels in the parasite, one can hypothesise that a decrease in redox-active ferrous heme in artemisinin resistant parasites leads to elevated GSH within the cell, potentially making them more tolerant to artemisinin-induced oxidative stress. However, this relationship, and its contribution to artemisinin resistance, is yet unstudied.

In this study we have addressed literature gaps around the mechanism of PfK13-mediated artemisinin resistance. We used state-of-the-art proteomics techniques and showed that PfK13 is the only protein that is consistently less abundant in several artemisinin-resistant clinical isolates in both ring-and trophozoite-stage parasites. Hemoglobin uptake and catabolism was then demonstrated to be decreased using heme fractionation assays and covalent activity-based probes. We further provide the first direct evidence for impaired artemisinin activation in PfK13-mutant artemisinin resistant parasites. This study also determined the role of GSH in PfK13-mediated resistance and revealed 1) that modulation of PfK13 levels or localisation modifies GSH levels, and (2) that elevated GSH decreases artemisinin potency using *in vitro* ring-stage survival assays. We then identified redox-modifying drugs that can potentiate artemisinin activity both *in vitro* and *in vivo*, and showed that sulforaphane (SFN), a common nutraceutical derived from broccoli sprout, can potentiate artemisinin activity against artemisinin resistant parasites both *in vitro* and *in vivo*.

## Study design

The study employs a multifaceted design combining *in vitro* and *in vivo* experiments to investigate artemisinin resistance mechanisms and potential interventions. Key elements of the study design include data-independent acquisition (DIA)-proteomics analysis performed on synchronized ring-and trophozoite-stages of artemisinin resistant and sensitive parasites to evaluate differential protein expression, with a focus on PfK13 protein levels. Hemoglobin, heme, and hemozoin were fractionated from artemisinin resistant and sensitive parasites, and absorbance was measured to assess the effects of artemisinin resistance on hemoglobin degradation products. Targeted analysis of thiol species was performed through LC-MS after extraction and derivatization of artemisinin resistant and sensitive parasite samples, with a focus on the GSH pathway. Streptolysin O-enriched ring-stage artemisinin resistant and sensitive parasites were exposed to DHA or arterolane, and activation of the drug was measured using LC-MS. Synchronised artemisinin resistant and sensitive parasites were treated with DHA or a combination of DHA with redox modulators (BSO, NAC and SFN) to evaluate their potentiating effect on DHA activity. Mice infected with *P. berghei* strains (artemisinin resistant and sensitive) were treated with artesunate or a combination of artesunate and SFN to evaluate therapeutic efficacy. This study integrates molecular, biochemical, and pharmacological approaches to elucidate artemisinin resistance mechanisms and explore redox modulators as a strategy to restore artemisinin sensitivity.

## Statistical analysis

In general, all data points are shown in the respective figures. Briefly, figure legends and the results section provide the descriptive statistics, which included mean, standard deviation, and range for each assay and experiment. For comparative analysis a T-test or ANOVA was used to compare the means of different groups (e.g., resistant vs. sensitive parasites) and determine if there were significant differences in protein abundance, thiol levels, or drug efficacy. Statistical analyses and visualization of the data was performed using GraphPad Prism.

## Parasite culture

*P. falciparum* parasites were cultured continuously in O+ human erythrocytes sourced from Australian Red Cross LifeBlood, with a hematocrit level maintained between 2% and 4%, following established protocols with slight modifications^9, 26^. Cultures were incubated at a constant temperature of 37 °C in an atmosphere comprising 94% N_2_, 5% CO_2_, and 1% O_2_. The culture medium used was RPMI 1640, enriched with 0.5% Albumax II (Gibco, Australia), HEPES buffer (5.94 g/L), hypoxanthine (50 mg/L), and sodium bicarbonate (2.1 g/L). To achieve parasite synchronization, cultures underwent multiple rounds of sorbitol lysis^27^. Parasite development was regularly assessed every 24-48 h through the examination of methanol-fixed and Giemsa-stained blood smears.

## IC_50_ and Ring stage survival assays

Ring stage survival assays were performed as previously described, with minor modifications^28^. Briefly, cultures were synchronised by multiple rounds of sorbitol treatment and when at the segmented schizont stage, parasites were magnet harvested and incubated for 3 h with fresh red blood cells to allow merozoite invasion. The cultures were again treated with sorbitol to eliminate remaining schizonts. The 0-3 hpi rings were adjusted to 1% parasitemia and 2% hematocrit and exposed to 700 nM of DHA for 1-6 h. After 1-6 h, the drug was removed by washing the cultures in complete RPMI culture medium (4 x 200 µL). Parasites were then returned to standard culture conditions for an additional 68 h. While for the IC_50_ assay, cultures adjusted to 1% parasitemia and 2% hematocrit were exposed to increasing concentrations of the inhibitor, with no drug wash. Vehicle-treated (DMSO) and unwashed (lethal DHA concentration for 72 h) parasites served as 100% and 0% growth controls, respectively. Parasite growth was assessed using the SYBR green I assay. Assays were performed in biological replicates (n= 2-9).

## Ring stage enrichment with Streptolysin O

Enriched ring-stage cultures were generated with Streptolysin O (SLO) as previously described with modifications^29^. Briefly, ring-stage cultures were tightly synchronised as described for ring stage survival assays and treated with a defined quantity of SLO units for 10-20 min at 37 °C to preferentially lyse uninfected erythrocytes. SLO activity was determined for each batch based on hemolytic activity, where one unit is defined as the amount of SLO required to lyse 50% of uninfected erythrocytes at 2% hematocrit in PBS for 30 min at 37 °C, as determined by titration. The cells were extensively washed 5-10 times with 50 mL of PBS to remove residual SLO and lysed RBC material. The resulting enriched ring-infected erythrocytes were resuspended in complete culture medium and used for subsequent peroxide activation and thiol metabolomics experiments.

## DIA-proteomics sample prep and analysis

### Sample preparation

The preparation of samples for untargeted proteomics was based on the method described in Siddiqui *et al.* (2022)^30^, with minor modifications. Samples each containing approximately 500 μg of protein in SDC lysis buffer (100 mM 4-(2-hydroxyethyl)-1-piperazineethanesulfonic acid [HEPES], 1% sodium deoxycholate [SDC], pH 8.1) were reduced and alkylated using 10 mM tris(2-carboxyethyl)phosphine (TCEP – Sigma Aldrich) and 40 mM of chloroacetamide (CAA) at 95 °C for 5 min. Proteins were digested into peptides overnight with trypsin (1:50; Promega, Australia), following digestions, samples were desalted using in-house-generated StageTips as described previously^31^. Eluted peptides were dried completely and reconstituted in 12 μL of loading buffer (2% acetonitrile in 0.1% trifluoroacetic acid), supplemented with indexed retention time (iRT) peptides (Biognosys, GmbH, Switzerland).

### Mass spectrometric instrumentation and data acquisition

LC-MS/MS was carried out as described previously^30, 32^, with minor modifications. Briefly, samples were loaded at a flow rate of 15 μL/min onto a reversed-phase trap column (100 μm x 2 cm), Acclaim PepMap media (Dionex), which was maintained at a temperature of 40 °C. The HPLC gradient was set to 158 min using a gradient that reached 30% of ACN after 123 min, then 34% of ACN after 126 min, 79.2% of ACN after 131 min and 2% after 138 min for a further 20 min. For data acquisition, a 25-fixed-window set-up of 24 *m/z* effective precursor isolation over the *m/z* range of 376–967 Da was applied. Full scan was performed at 70,000 resolution (AGC target of 2e^5^, maximum injection time set to auto, normalized collision energy of 27.0).

### Proteomics data analysis

The proteomics data were processed using Spectronaut™ software (version 13.0), referencing an in-house–generated *Plasmodium falciparum* spectral library as previously described^30^ and was based on default Spectronaut™ settings (Manual for Spectronaut™ 13.0, available on Biognosis website). Quantification was performed at the fragment level with channel interference removal enabled, a 1% false discovery rate was used for protein, peptide and peptide spectral match level identification, inbuilt normalisation was applied with a minimum three fragment ions used for quantification. Tryptic digestion was used with specific cleavage rules and a missed cleavage tolerance of two. Carbamidomethylation of cysteines was set to a fixed modification while acetylation of N-termini and oxidation of methionine were set as variable. Protein level abundance values were imported into the Monash Proteomics Analyst Suites DIA-Analyst tool (v.0.8.5) for statistical analysis and visualisation. Differential proteins were determined using a p-value of ≤0.05 and a log2 fold-change of 0.5. The mass spectrometry proteomics data have been deposited to the ProteomeXchange Consortium via the PRIDE^33^ partner repository with the dataset identifier PXD068108.

### Hemoglobin fractionation assay

The hemoglobin fractionation assay was adapted from published methods^32, 34^ with minor modifications. Hemoglobin, heme, and hemozoin fractions were isolated from 3-6 hpi ring-stage parasite pellets using 20 μL of the previously defined reagents^32, 34^ (or 50 μL for trophozoite stages). The absorbance of each fraction was measured at a wavelength of 405 nm using a Perkin Elmer Ensight Plate Reader. Statistical analysis and visualisation of results used GraphPad Prism 9.1.0 software.

### Measurement of cysteine protease activity using covalent activity-based probes

The activity-based probe, FY01, containing a Cy5 fluorophore, was used to measure cysteine protease activity in artemisinin resistant and sensitive strains as previously described^35, 36^. Briefly, tightly synchronised 22-24 h artemisinin resistant and sensitive trophozoite-stage parasites were purified by lysing the red blood cells using 0.1% saponin on ice. Parasite pellets were then lysed by sonication in citrate buffer (50 mM trisodium citrate [pH 5.5], 0.5% CHAPS, 0.1% Triton X-100, 4 mM dithiothreitol) or phosphate buffered saline (PBS) (pH 7.2). Protein concentration was determined using BCA protein assay (Pierce) and an equal amount of each sample was incubated with FY01 at 1 μM for 30 min at 37 °C to label active cysteine proteases. The reaction was then quenched by the addition of 5x reducing buffer (50% glycerol, 250 mM Tris-Cl [pH 6.9], 10% SDS, 0.05% bromophenol blue, 6.25% beta-mercaptoethanol), boiled and separated by sodium dodecyl sulfate polyacrylamide (SDS-PAGE) on 15% polyacrylamide gels. Visualization was achieved by direct scanning of the gel for Cy5 fluorescence. Images were processed and quantified in ImageJ 1.51f.

### Targeted thiol analysis

#### Sample preparation

Thiol concentrations within *P. falciparum*, particularly focusing on the GSH pathway, were quantified using a targeted liquid chromatography-mass spectrometry (LC-MS) approach. The method of preparing targeted thiol samples adhered to previously established protocols^4^. SLO-enriched (3-6 hpi ring-stage) and magnet-enriched (22-24 hpi trophozoite-stage) parasites at 3-5 x 10^7^ cells were quenched and extracted using freshly prepared N-Ethylmaleimide (NEM) extraction solvent. This solvent was composed of 40 µL of 50 mM NEM (Sigma-Aldrich) in a mixture of 80% methanol and 20% 10 mM ammonium formate (Sigma-Aldrich), combined with 40 µL of 100% acetonitrile (Thermo Fisher). The mixture was then agitated at 4 °C for one hour and subsequently centrifuged at maximum speed for 10 min. Following this, 75 µL of the metabolite extract was transferred into LC-MS glass vials and stored at −80 °C until the time of analysis.

### Liquid chromatography–mass spectrometry analysis and data processing

Derivatized thiols were analysed as described previously^4^. To minimize variability, samples were analysed in a single batch and randomized to mitigate potential LC-MS system drift effects over the duration of the analysis. Data processing involved the quantification of metabolites through the integration of MS1 peaks using QuanBrowser software (Thermo Scientific), adhering to a predefined protocol^4^. Metabolite identification was accomplished by comparing the accurate mass and retention time of NEM-derivatized samples against those of authentic reference standards. Quantitative data, represented by the calculated peak areas, were visualized using GraphPad Prism software version 9.1.0. Statistical analysis to determine significant differences between groups was performed using GraphPad Prism software.

### Peroxide activation and LC-MS analysis

Peroxide drug activation was assessed in SLO-enriched (parasitemia used varied for different experiments and is stated in the figure legends) early ring-stage cultures (0-3 hpi) at 20% hematocrit. Uninfected RBCs at the same hematocrit from the same donor were analysed in parallel. Cultures were incubated with 700 nM of DHA or 1 µM of arterolane at 37 °C, followed by periodic sampling between 0 h and 2 h. At each sampling point, cellular Fe^2+^ was oxidised to Fe^3+^ by addition of potassium dichromate (40 mM final concentration) to prevent further peroxide bond activation, followed by protein precipitation with acetonitrile (containing 150 ng/mL of diazepam as the internal standard) as previously described^28^. The samples were transferred to microcentrifuge tubes, vortex mixed, and allowed to extract on ice for 10 min. After centrifugation for 10 min at 4 °C, the supernatant was transferred to analytical vials and stored at -80 °C until LC-MS analysis.

Samples were analysed using high-resolution mass spectrometry (Q Exactive, Thermo Fisher) coupled with a Dionex Ultimate 3000 ultra-high-performance liquid chromatography (UHPLC) system (Thermo Fisher) as previously described with some modifications^37^. Briefly, analytical separation was performed on a 100 mm by 2.1 mm, 2.7-mm Ascentis Express C8 reversed phase column, with a guard column of the same material (Sigma-Aldrich). Compounds were eluted using a binary gradient solvent system consisting of 0.1% formic acid (solvent A) and 0.1% formic acid in acetonitrile (solvent B). The gradient profile was as follows: 0 to 1 min, 10% B; 1 to 4.5 min, 25% B; 4.5 to 5 min 50% B; 5 to 8.5 min 95% B; 8.5 to 9.5 min, 95% B; 9.5 to 10 min, 10% B; and 10 to 12 min 10% B. Analytes were eluted at a flow rate of 0.3 mL/min and detected on the mass spectrometer in positive ionisation mode. Targeted analysis based on accurate mass and retention time, and integration of LC-MS peak areas for the analytes of interest, was performed using Thermo Xcalibur Quan Browser (version 4.2 SP1). Activation was determined by measuring the percentage of DHA or arterolane remaining as a function of time, with activation rates determined by log-linear regression assuming pseudo-first order degradation.

### Sulforaphane metabolomics sample preparation and analysis

Magnet harvested (>80% parasitaemia) Pf3D7 wildtype parasites were treated with 30 µM of sulforaphane (SFN) for 3 h or DMSO control. After drug treatment, metabolites of magnet harvested parasites were extracted with methanol as previously described^38^. Metabolite analysis was performed by liquid chromatography-mass spectrometry LC-MS using hydrophilic interaction liquid chromatography (HILIC) and high-resolution (Q-Exactive Orbitrap, Thermo Fisher) MS as previously described^32, 38^. Briefly, samples (10 μL) were injected onto a Dionex RSLC U3000 LC system (Thermo) fitted with a ZIC-pHILIC column (5 μm particle size, 4.6 by 150 mm; Merck) and 20 mM ammonium carbonate (A) and acetonitrile (B) were used as the mobile phases. A 22 min gradient starting from 80% B to 50% B over 15 mins, followed by washing at 5% B for 3 mins and re-equilibration at 80% B, was used. MS with a heated electrospray source was operating in positive and negative modes (rapid switching) and a mass resolution of 35,000 from *m/z* 85 to 1275. Sample injections within the experiment were randomized to avoid any impact of systematic instrument drift on metabolite signals. Retention times for ∼350 authentic standards were checked manually to aid metabolite identification.

Metabolomics data sets were analysed using IDEOM^39^. Raw files were converted to mzXML with msconvert^40^, extraction of LC-MS peak signals was conducted with the Centwave algorithm in XCMS, alignment of samples and filtering of artifacts with mzMatch, and additional data filtering and metabolite identification was performed in IDEOM (Supp Data Sheet 3). All metabolite identifications are based on accurate mass and retention time (or predicted retention time where standards were not available), and should be considered putative, as features may represent alternative isomers. Metabolite abundance of 1110 putative metabolites was determined by LC-MS peak height. Statistical analyses used t test (p-value< 0.05) and Volcano plot generated using IDEOM^39^. Raw metabolomic data is available at the NIH Common Fund’s National Metabolomics Data Repository (NMDR) website, the Metabolomics Workbench, https://www.metabolomicsworkbench.org, where it has been assigned Project ID (ST004194).

### Sulforaphane artesunate *in vivo* experiments

Female ARC(s) Swiss mice (6-10 weeks) were sourced from the Animal Resource Centre (Perth, Australia) and maintained on a standard rodent diet and housed under controlled conditions (21°C, 12:12 hour light: dark cycle). All experiments were performed in accordance with the recommendations of the National Health and Medical Research Council Australian Code of practice for the care and use of animals for scientific purposes, and approved by the Deakin University Animal Welfare Committee (Project G11-2023). For *in vivo* studies, *Plasmodium berghei* ANKA clone 1804cl1 artesunate-sensitive parasites were used (1804^wt^) together with parasites containing a PfK13 mutation (G1989; 1804^M488I^) on the same background^41^. Infections in donor mice were established by intraperitoneal (ip) injections of 100-200 µL of cryopreserved parasite stocks and parasite load monitored via Giemsa-stained blood smears. Cardiac bleeds were performed when parasite loads were >2 %. Experimental mice in cohorts of 3-6 were infected ip on day 0 with between 1 × 10^6^ and 1 × 10^7^ iRBC harvested from donor mice. Once a parasitemia of ∼2% was reached, groups of mice were injected ip daily for 3 days with artesunate (Sigma; 30-60 mg/kg diluted in DMSO) or artesunate with SFN (Sigma; 5-20 mg/kg) and parasite burden was monitored daily by Giemsa stain. Cardiac bleeds were performed when mice reached a parasitemia of >15 % or were symptomatic.

## Results

### Proteomic analysis of artemisinin resistant and sensitive 3-6 hpi ring-stage parasites demonstrates altered abundance of PfK13

A number of studies, including from our own lab, have performed comparative proteomic analysis of trophozoite stage PfK13-mediated artemisinin resistant and sensitive South East Asian *P. falciparum* isolates^9, 11, 30^. Here, we have used a DIA mass spectrometry approach to compare a range of artemisinin resistant (Cam3.II^R539T^, Cam3.II^C580Y^ and MRA1240) and sensitive parasites (Cam3.II^REV^ and MRA1239) and for the first time also performed comparative proteomics analysis in 3-6 hpi ring-stage parasites, which is the stage at which artemisinin resistant parasites display differential survival relative to sensitive parasites (Supp data sheet 1). Across Cam3.II^R539T^ and Cam3.II^C580Y^, compared to their isogenic sensitive isolate Cam3.II^REV^ (isolates from Pursat Province, Cambodia) in both 3-6 hpi ring-(Fig 1) and in 24 hpi trophozoite-stage parasites (previously published)^9^, we found three proteins to be consistently dysregulated in abundance. These included an ATP-dependent RNA helicase (PF3D7_0320800), *Plasmodium* exported protein (PHISTb – PF3D7_0401800) and PfK13 itself, all of which were significantly decreased in resistant compared to sensitive parasites (Fig 1A and B, Supp Table 1). In addition, in early ring-stage parasites, PfK13 mutation down-regulated the abundance of several proteins involved in the parasite respiratory chain (Supp Table 1). Comparative proteomics analysis of 24 hpi trophozoite-stage MRA1240 (R539T PfK13 mutation) and MRA1239 (artemisinin sensitive isolate) isolates from Western Cambodia showed that the only consistently significantly dysregulated protein was PfK13 (Fig 1C, Supp Fig 1 and Supp data sheet 2). PfK13 levels were consistently and significantly down-regulated by on average 1.5-fold across both ring-and trophozoite-stage parasites independent of PfK13 mutation and resistant isolate (Fig 1C).

**Figure 1.**
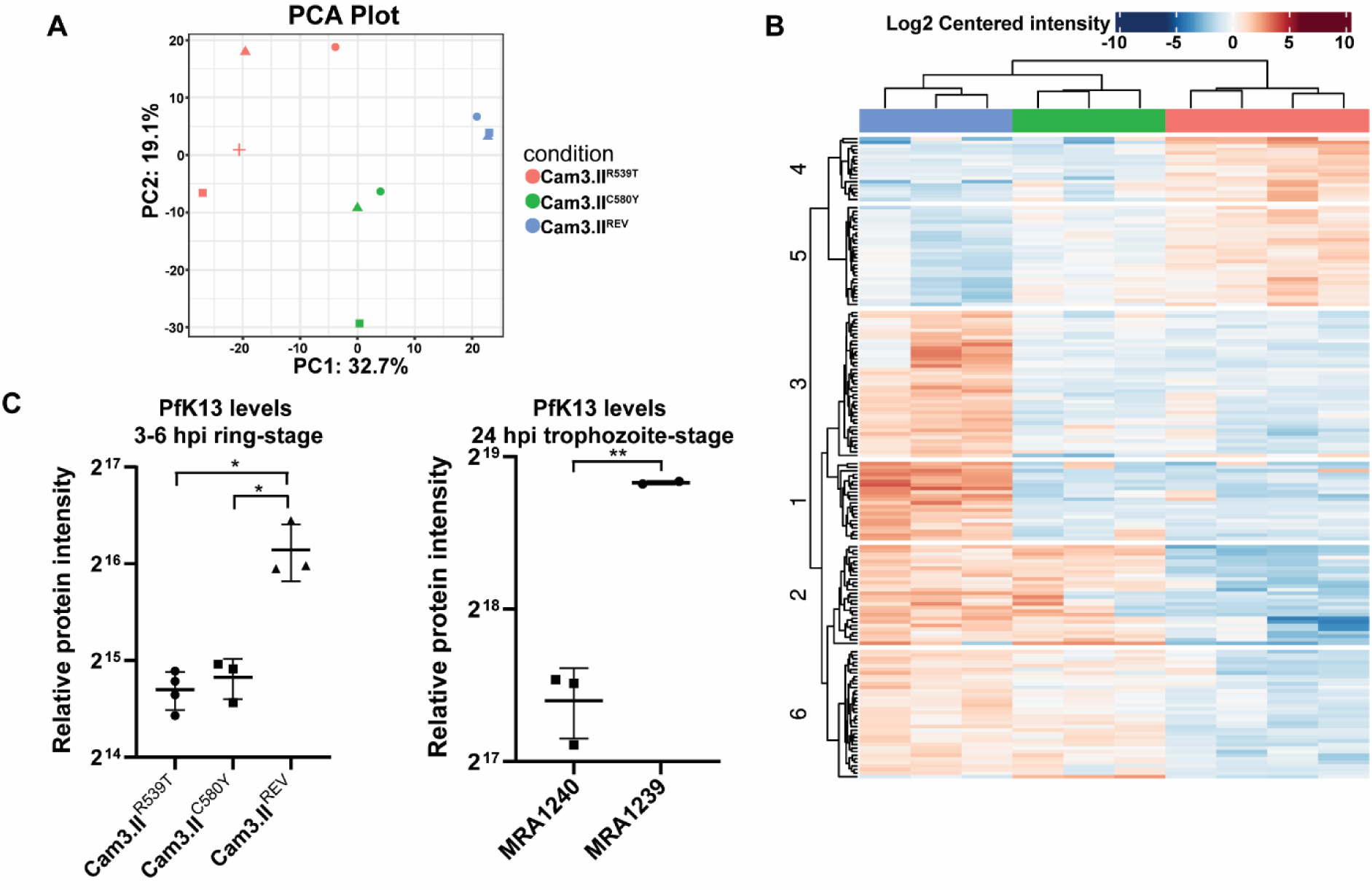
Proteomics analysis of artemisinin resistant and sensitive parasites. (**A**) Principal component analysis of all the 1622 proteins quantified across 3-6 hpi ring stage Cam3.II^R539T^ (artemisinin resistant, 4 samples), Cam3.II^C580Y^ (artemisinin resistant, 3 samples) and Cam3.II^REV^ (artemisinin sensitive, 3 samples). (**B**) Hierarchical clustering analysis of significantly dysregulated proteins across the conditions (blue – Cam3.II^REV^, green – Cam3.II^C580Y^ and pink – Cam3.II^R539T^) (log2 fold-change of 0.5 and Bonferroni corrected P-value of less than 0.05). (**C**) Relative abundance of PfK13 in 3-6 hpi ring-stage and 24 hpi trophozoite-stage artemisinin resistant and sensitive parasites. Cam3.II^R539T^, Cam3.IIC^580Y^ and MRA1240 (artemisinin resistant parasites). Cam3.II^REV^ and MRA1239 (artemisinin sensitive parasites). Values represent the means of two-four independent experiments ± the standard errors of the mean (SEM), p-value was calculated using Welch’s t-test (*, p < 0.05; **, p < 0.01;***, p-value < 0.001).

### Hemoglobin digestion is decreased in artemisinin resistant parasites

PfK13 has been localised to the parasite cytostome and shown to be involved in hemoglobin uptake. Thus, decreased PfK13 protein abundance is likely to be associated with impaired hemoglobin uptake and digestion^7–10^. It is therefore hypothesised that this decreases the availability of free heme to activate artemisinin, however, this is yet to be demonstrated in early ring-stage clinical isolates. Here, we have used a modified hemoglobin fractionation assay^32, 34^ in 3-6 hpi ring-and 24 hpi trophozoite-stage artemisinin resistant (Cam3.II^R539T^, Cam3.II^C580Y^ and MRA1240) and sensitive (Cam3.II^REV^ and MRA1239) parasites to measure hemoglobin, heme and hemozoin species. We show that in 3-6 hpi ring-stage parasites, heme levels were significantly decreased in abundance in resistant parasites compared to sensitive parasites, while hemoglobin and hemozoin levels were not significantly different (Fig 2A). In 24 hpi trophozoite-stage parasites, hemoglobin and heme levels were not measurably different, but hemozoin levels were significantly decreased in resistant parasites compared to sensitive parasites (Fig 2B), indicative of decreased turnover of heme into hemozoin in resistant parasites over the preceding 24 h of the parasite lifecycle.

**Figure 2.**
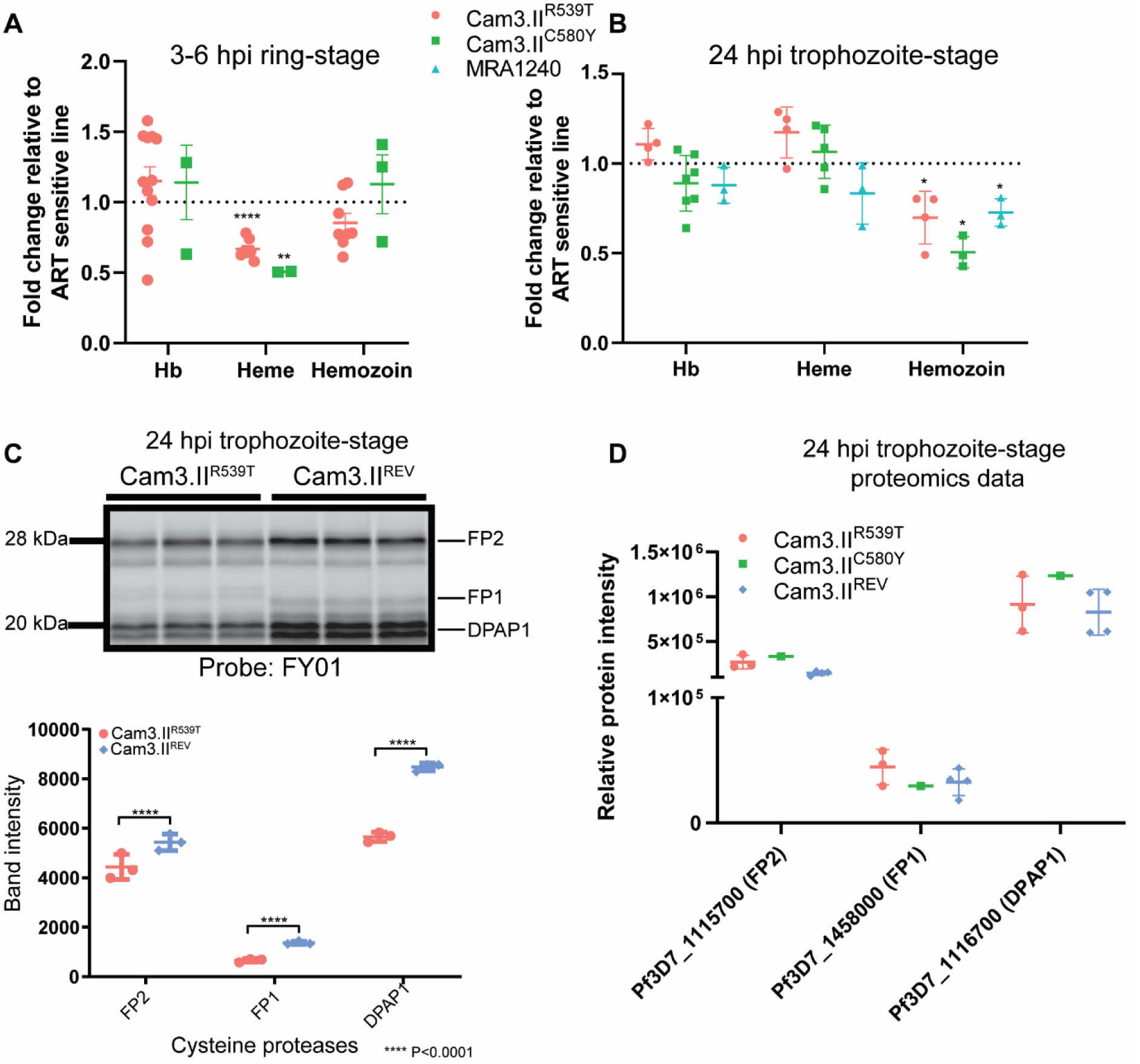
Hemoglobin digestion is perturbed in artemisinin resistant parasites. Hemoglobin (Hb) and its degradation products (heme and hemozoin) fold-change in 3-6 hpi ring-stage (**A**) and 24 hpi trophozoite-stage (**B**) of artemisinin-resistant compared to artemisinin-sensitive *P. falciparum*. Values represent the means of two-ten independent experiments ± the standard errors of the mean (SEM). p-value was calculated using one sample t-test with a hypothetical mean comparison of 1 (*, p < 0.05; **, p < 0.01, ****, p<0.0001). (**C**) Parasite cysteine protease activity in artemisinin resistant (Cam3.II^R539T^) and sensitive (Cam3.II^REV^) parasites using the activity-based probe, FY01. Cysteine protease activity and densitometric analysis of the falcipain 1 (FP1), falcipain 2/3 (FP2) and dipeptidyl aminopeptidase 1 (DPAP1) signal in 24 hpi trophozoite-stage artemisinin resistant and sensitive *P. falciparum*. FY01 labelling was detected by scanning of in-gel Cy5 fluorescence. The lanes are independent biological replicates. For the densitometric analysis, the FP1, FP2 and DPAP1 signal intensity was normalised to the average signal intensity of the appropriate time point in the untreated control (±SD). p-value was calculated using Welch’s t-test (****, p-value < 0.0001). (**D**) Relative abundance of FP1, FP2 and DPAP1 from 24 hpi trophozoite-stage proteomics analysis of artemisinin resistant (Cam3.II^R539T^ (3 replicates) and Cam3.II^C580Y^ (1 replicate)) and sensitive (Cam3.II^REV^ (4 replicates)) *P. falciparum*, data taken from Siddiqui *et al.* 2022^30^.

To assess whether a decrease in hemoglobin uptake is also associated with perturbed hemoglobin digestion, we used covalent activity-based probes^35, 36, 42^ to measure the activity of hemoglobin-digesting enzymes in artemisinin resistant (Cam3.II^R539T^) and sensitive (Cam3.II^REV^) parasites. We showed that the activity of cysteine proteases (falcipain2 and 3, and dipeptidyl aminopeptidase 1)^43^ involved in hemoglobin digestion were significantly decreased in 24 hpi artemisinin resistant parasites compared to sensitive (Fig 2C), while proteomics analysis showed no difference in the abundance of these proteins (Fig 2D)^30^. We propose that the activity of these cysteine proteases is dependent on the pH and protease activity of a fully functional digestive vacuole, and that normal DV formation is impaired in K13-mutant parasites^7^.

### Artemisinin activation is decreased in PfK13 mutant lines

Artemisinin requires activation by free heme, thus we anticipated that the decrease in heme levels observed in 3-6 hpi ring-stage parasites would result in decreased artemisinin activation and, therefore, decreased activity. Here, we used LC-MS to measure the activation rate of dihydroartemisinin (DHA) or the related peroxide antimalarial, arterolane. Test compounds were incubated in enriched 3-6 hpi ring-stage artemisinin resistant (Cam3.II^R539T^, Cam3.II^C580Y^ and MRA1240) or sensitive (Cam3.II^REV^ and MRA1239) parasites and the level of peroxide activation determined by measuring the percentage of intact compound remaining as a function of time (Fig 3A, B and D). Elevated levels of intact peroxide antimalarials were observed in resistant parasites over the duration of the incubation, corresponding to a slower rate of activation compared to sensitive parasites. On average (dependent on the parasitemia of each individual experiment), the percentage of DHA remaining inactivated in artemisinin resistant parasites (Cam3.II^R539T^ and MRA1240) following a 2 h incubation was significantly higher (2-fold greater) than sensitive (Cam3.II^REV^ and MRA1239) (Fig 3F).

**Figure 3.**
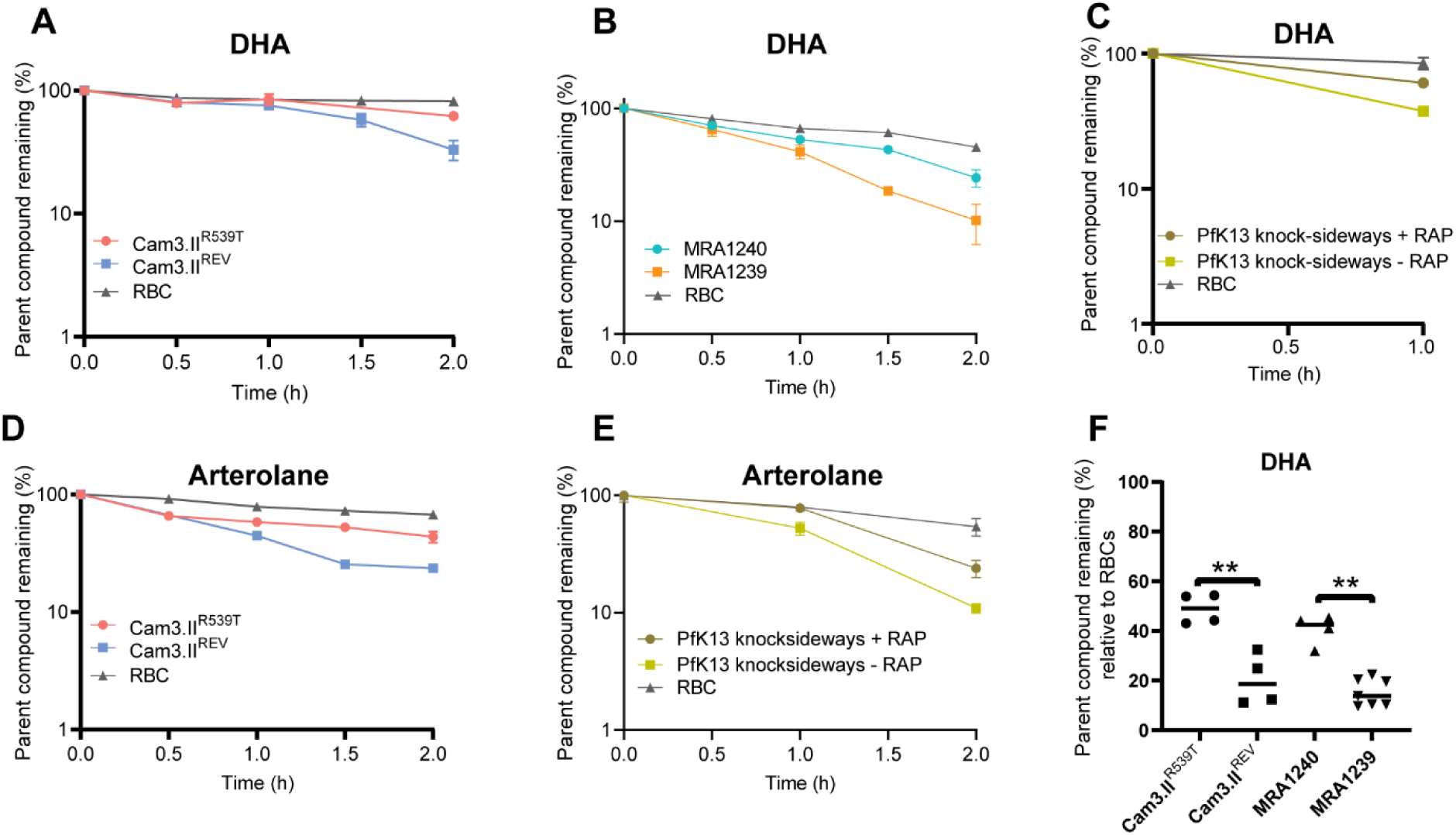
Peroxide activation is perturbed in artemisinin resistant and PfK13 knock-sideways parasites. Dihydroartemisinin (DHA) (**A**, **B** & **C**) and arterolane (**D** & **E**) degradation profiles in 3-6 hpi ring-stage artemisinin resistant (Cam3.II^R539T^ (red) and MRA1240 (teal)), artemisinin sensitive (Cam3.II^REV^ (blue) and MRA1239 (orange)), PfK13 knock-sideways parasites with (brown) or without (yellow) rapamycin (RAP) parasites and in uninfected red blood cells (gray). The starting concentration of DHA was 700 nM and arterolane was 1 µM. Values represent the means from a single biological experiment ± the standard deviation of technical replicates. (**F**) Percentage DHA remaining relative to uninfected red blood cells control and normalized to parasitemia across two biological replicates for artemisinin resistant (Cam3.II^R539T^ and MRA1240) and sensitive (Cam3.II^REV^ and MRA1239) parasites. p-value was calculated using paired t-test (**, p < 0.01).

We also demonstrated that this slower rate of activation is directly linked to PfK13 levels at the parasite periphery, by analysis of a PfK13 knock-sideways parasite line^8^. These parasites, in the presence of rapamycin (RAP) mislocalize PfK13 away from its native subcellular location and into the nucleus, thereby limiting its native function. We showed that the rate of peroxide antimalarial activation is significantly decreased in PfK13 knock-sideways parasites in the presence of RAP compared to the same parasites without any RAP (Fig 3C and E). Collectively, these results directly demonstrate that resistance-causing PfK13 mutations in early rings result in a 1.5-fold loss in PfK13 protein abundance, which decreases intra-parasitic heme levels and subsequently impairs peroxide antimalarial activation.

### Artemisinin resistant parasites have increased thiol levels

Our previous metabolomics studies suggested that the cellular environment of artemisinin resistant parasites is in a more reduced state, whereby the level of reduced glutathione (GSH) was significantly elevated in 24 hpi trophozoite-stage resistant parasites compared to sensitive. Here, we have used our quantitative thiol-metabolomics method to more accurately measure the levels of derivatised thiol species in a range of artemisinin resistant (Cam3.II^R539T^, Cam3.II^C580Y^, PL7, and MRA1240) and sensitive (Cam3.II^REV^, PL2 and MRA1239) isolates in both 3-6 hpi ring-and 24 hpi trophozoite-stage parasites (Fig 4). PL2 encoding the wildtype PfK13 and PL7 encoding the PfK13 mutant (R539T) are another set of field isolates from Cambodia. Elevated levels of reduced GSH and its precursor γ-glutamyl cysteine (gGlu-Cys) were observed in resistant parasites, while oxidised glutathione (GSSG) was lower (Fig 4A, B & C). We also determined thiol levels in the PfK13 knocksideways line in the presence of RAP and found that mislocalisation of PfK13 significantly elevated levels of reduced GSH and its precursor gGlu-Cys (Fig 4D). This was not due to RAP treatment, as thiol levels were not altered by RAP treatment in wildtype Pf3D7 parasites (Fig 4D). Together, this suggests that altered PfK13 modulates the levels of reduced GSH, leading to a more reducing cellular environment in artemisinin-resistant parasites.

**Figure 4.**
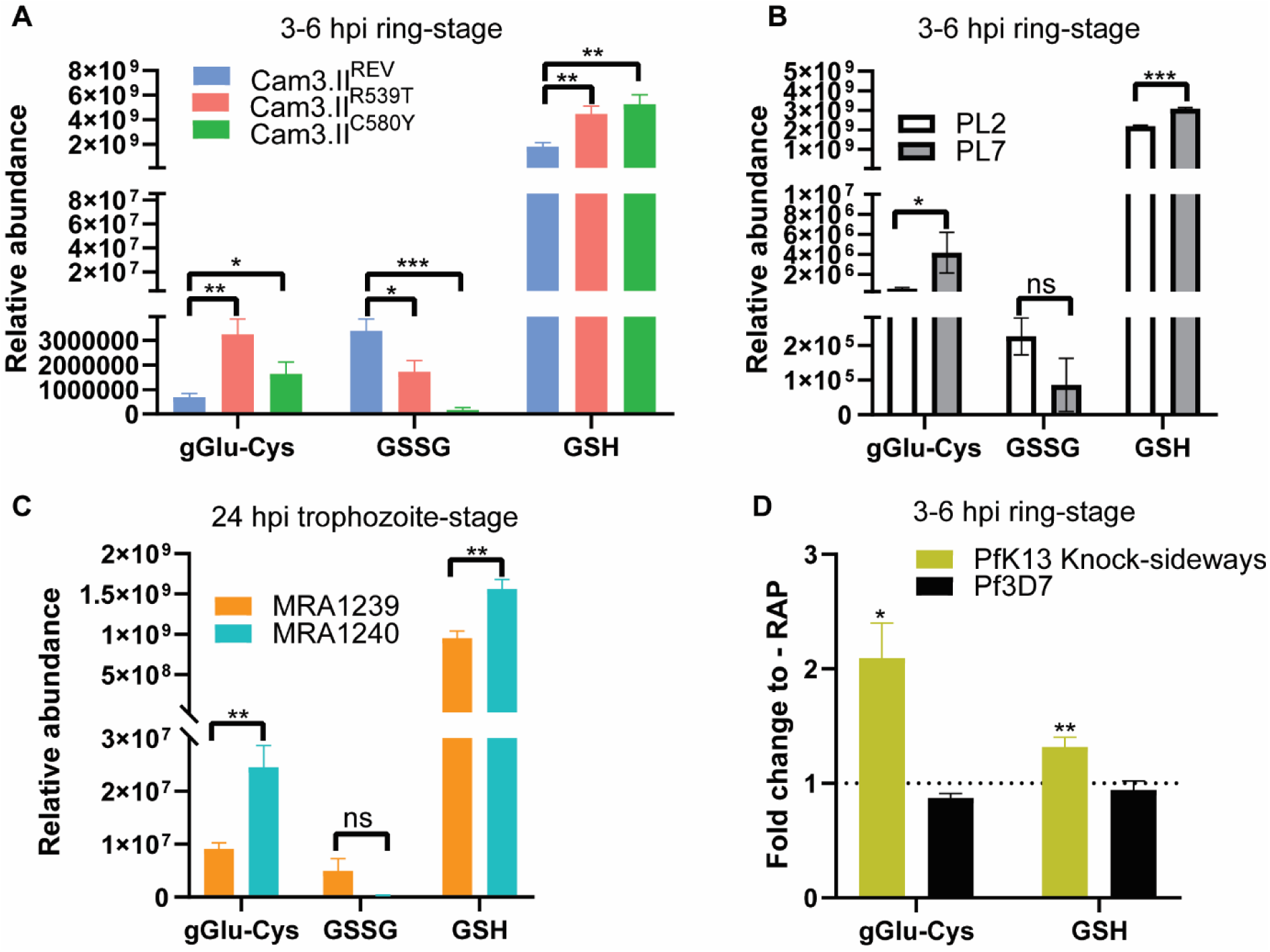
Thiol levels are dysregulated in artemisinin resistant parasites. (**A** & **B**) Relative levels of NEM-derivatized γ-glutamyl cysteine (gGlu-Cys), oxidised glutathione (GSSG), reduced glutathione (GSH) in 3-6 hpi ring-stage artemisinin resistant (Cam3.II^R539T^, Cam3.II^C580Y^, PL7) and sensitive parasites (Cam3.II^REV^ and PL2) and (**C**) 24 hpi trophozoite stage artemisinin resistant (MRA1240) and sensitive (MRA1239) parasites. p-value was calculated using Welch’s t-test (*, p < 0.05; **, p < 0.01;***, p-value < 0.001). (**D**) NEM-derivatised gGlu-Cys and GSH fold change in PfK13 knock-sideways and Pf3D7 lines following treatment with rapamycin (RAP). Thiol measurement is from three to five biological replicates. p-value was calculated using one sample t-test with a hypothetical mean comparison of 1 (*, p < 0.05; **, p < 0.01).

In order to investigate whether elevated levels of reduced GSH were due to less redox active heme iron being liberated from impaired hemoglobin uptake and digestion in resistant parasites, 24 hpi trophozoite-stage parasites were treated for 3 h with 10 µM of the cysteine protease inhibitor, E64d, to block hemoglobin digestion and decrease free heme levels. The hemoglobin fractionation assay confirmed that E64d-treated parasites have significantly decreased heme and hemozoin levels, while undigested hemoglobin levels were elevated following treatment (Supp Fig 2A). We then determined thiol levels and, surprisingly, thiol levels in E64d-treated parasites were unaffected (Supp Fig 2B). This suggests that levels of free heme alone are not sufficient to immediately change the oxidative state of the parasite, although it is recognised that chemical inhibition of cysteine proteases may have additional impacts on the parasite, and does not perfectly mimic the heme depletion caused by K13-mediated alteration of hemoglobin uptake.

### Targeting parasite oxidative capacity can re-sensitise resistant parasites

Artemisinin resistant parasites have a more reduced oxidative status and we have previously shown that artemisinin disproportionately alkylates proteins involved in parasite redox homeostasis and that disrupted redox processes are involved in the artemisinin mechanism of action^4^. Therefore, we hypothesised that targeting parasite redox processes (more specifically, decreasing levels of reduced GSH) using available redox modulators, would re-sensitise resistant parasites to artemisinin. We first used a known γ-glutamyl cysteine synthetase inhibitor, buthionine sulfoximine (BSO)^44^. γ-Glutamyl cysteine synthetase is the rate-limiting enzyme in the *de novo* biosynthesis of GSH from cysteine, glutamate and glycine^23^.

Treatment of early ring-stage artemisinin resistant (Cam3.II^R539T^) and sensitive (Cam3.II^REV^) parasites with 10 mM of BSO for 1 h significantly depleted both γ-Glu-Cys and reduced GSH levels in both lines compared to untreated controls (Fig 5A). This confirms that BSO inhibits parasite γ-glutamyl cysteine synthetase in *P. falciparum*. We then pre-incubated early ring-stage artemisinin resistant and sensitive parasites with BSO for 1 h prior to pulse treatment with 700 nM of DHA for 6 h and measured survival, analogous to the standard *in vitro* Ring-stage Survival Assay (RSA) for artemisinin resistance. Pre-treatment with BSO hypersensitised parasites to DHA treatment, while BSO alone at the same dose and duration did not affect parasite growth (Fig 5B). This demonstrates that BSO potentiates artemisinin activity in both sensitive and resistant parasites.

**Figure 5.**
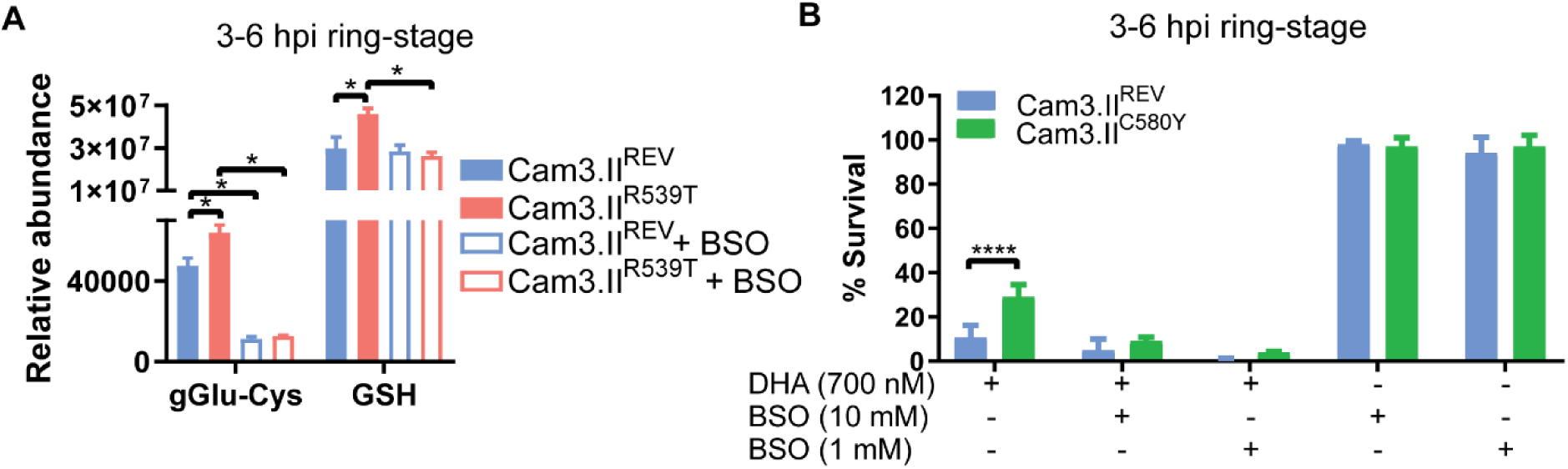
Decreasing glutathione levels can increase artemisinin sensitivity. (**A**) Relative levels of NEM-derivatized γ-glutamyl cysteine (gGlu-Cys), and reduced glutathione (GSH) in 3-6 hpi ring-stage artemisinin resistant (Cam3.II^R539T^ (red)) and sensitive parasites (Cam3.II^REV^ (blue)) following treatment (clear bars) with 10 mM of buthionine sulfoximine (BSO) for 1 h. Thiol measurement is from three biological replicates. p-value was calculated using Welch’s t-test (*, p < 0.05). (**B**) 3-6 hpi Ring-stage survival assay determination for artemisinin resistant (Cam3.II^C580Y^ (green)) and sensitive (Cam3.II^REV^ (blue)) parasites following treatment with 700 nM of DHA for 6 h alone, DHA with 10 mM of BSO (pre-treated for 1 h), DHA with 1 mM BSO (pre-treated for 4 h), and BSO alone at 10 mM and 1 mM. Values represent the means of six independent experiments ± the standard errors of the mean (SEM), p-value was calculated using Welch’s t-test (****, p-value < 0.0001).

In addition to this, we had previously generated a PfDd2-derived parasite line with a copy number amplification (2-3-fold) of a region in chromosome 1, which included 36 genes including *pfmrp1* (*Plasmodium falciparum* multidrug resistance protein 1; PfMRP1)^45^ (Supp Fig 3A). PfMRP1 has been hypothesised to be involved in drug efflux, but also efflux of reduced GSH. Therefore, we first measured the levels of thiol species in 3-6 hpi ring-stage PfMRP1-mutant parasites and compared to its wildtype parental control (PfDd2). Cellular levels of both γ-Glu-Cys and reduced GSH levels were significantly decreased (Supp Fig 3B), consistent with our hypothesis that the PfMRP1 mutant increases efflux of reduced GSH. Then, to determine whether the decreased levels of reduced GSH would affect sensitivity to artemisinin treatment, PfMRP1-mutant and PfDd2 parasites were pulse treated with different concentrations of DHA (100 – 700 nM) for 1 h and the PfMRP1 mutant was significantly more sensitive to DHA treatment compared to PfDd2 wildtype (Supp Fig 3C). In order to ensure that these changes in DHA sensitivity were not due to altered heme levels, we measured heme levels in PfMRP1-mutant and DD2 wildtype parasites and found no differences (Supp Fig 3D). This provides additional evidence that decreased parasite thiol levels directly influence artemisinin sensitivity.

In order to determine the impact of elevated GSH on artemisinin potency, we pre-treated artemisinin resistant (Cam3.II^C580Y^) and sensitive (Cam3.II^REV^) parasites with N-acetyl cysteine (NAC), a precursor for GSH *de novo* synthesis^46^ and measured sensitivity to 700 nM of DHA by RSA. Decreased DHA potency was observed following pre-incubation with NAC (Fig 6A). Measurement of the rate of artemisinin activation showed that there was no significant difference in DHA activation following pre-treatment with NAC (Fig 6B). We also investigated the effect of NAC treatment on heme levels and found no significant differences (Fig 6C), thereby indicating that thiol elevation (rather than altered drug activation) is primarily responsible for the impact of NAC on artemisinin sensitivity.

**Figure 6.**
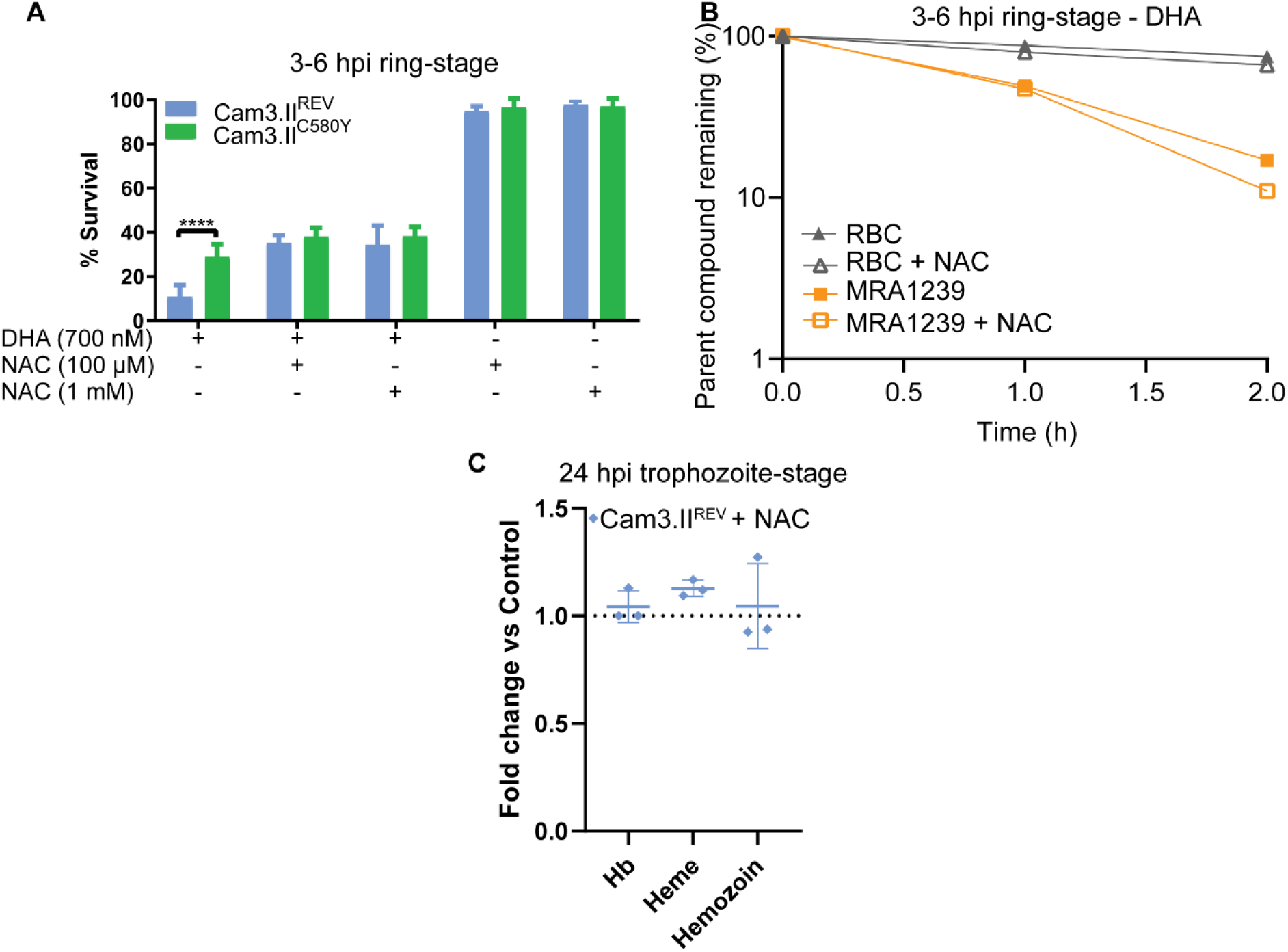
Increasing glutathione levels decreases artemisinin sensitivity. (**A**) 3-6 hpi Ring-stage survival assay determination for artemisinin resistant (Cam3.II^C580Y^ (green)) and sensitive (Cam3.II^REV^ (blue)) parasites following treatment with 700 nM of DHA for 6 h alone, DHA with 100 µM of N-acetyl cysteine (NAC) (pre-treated for 1 h), DHA with 1 mM NAC (pre-treated for 1 h), and NAC alone at 100 µM and 1 mM. Values represent the means of six independent experiments ± the standard errors of the mean (SEM), p-value was calculated using Welch’s t-test (****, p-value < 0.0001). (**B**) DHA (700 nM) degradation profiles in 3-6 hpi ring-stage artemisinin sensitive (MRA1239 (orange)) parasites and in uninfected red blood cells (gray) following pre-treatment with 1 mM NAC for 1 h. Values represent the means of technical replicates from one experiment ± the standard deviation of the mean (SD). (**C**) Hemoglobin (Hb) and its products (heme and hemozoin) fold-change in 24 hpi trophozoite-stage artemisinin-sensitive (Cam3.II^REV^) parasites treated with 1 mM NAC for 1 h. Values represent the means of three independent experiments ± the standard deviation of the mean (SD).

### *In vitro* data shows that sulforaphane (SFN) potentiates artemisinin activity

SFN is an active phytochemical found in *Brassica* vegetables, and is the hydrolytic product of glucosinolates metabolised by myrosinase. In mammalian cells, SFN conjugates GSH, either passively or through the activity of glutathione-S-transferases, and the SFN-GSH conjugate causes oxidative stress^47^. In response to this stress, the Kelch-like ECH-associated protein (Keap1) dissociates from the nuclear factor erythroid 2-related factor 2 (Nrf2). Nrf2 translocates to the nucleus and interacts with the antioxidant response element (ARE) of target genes, resulting in expression of antioxidant genes, which induces an antioxidant response shown to be beneficial for several human conditions^48^. However, *P. falciparum* has no identified Nrf2 orthologue and so likely lacks a KEAP1-Nrf2 mediated antioxidant response, which suggests that the SFN-GSH conjugate should only cause oxidative stress in parasites. We therefore hypothesised that SFN should have anti-parasitic activity, and determined an IC_50_ of 11.8 ± 0.4 µM for SFN in a standard growth inhibition assay in Pf3D7 wildtype parasites (Supp Fig 4A). After demonstrating the anti-parasitic effect *in vitro*, we performed untargeted metabolomics on parasites treated with 30 µM of SFN for 3 h, and the most significantly elevated metabolites in treated cells compared to controls were SFN-thiol adducts, including SFN-GSH and SFN-Cysteine. Interestingly, SFN-GSH was the 3rd most abundant metabolite detected in this study (after phosphorylcholine and PC(34:1)) based on LC-MS peak height, while SFN adducts with other peptides and amino acids were detected a low levels (Fig 7A and B). The most significantly depleted metabolites following SFN exposure were short peptides, which is consistent with the metabolite signature similar seen with other fast acting antimalarials such as DHA^36^. The metabolomics data confirms that SFN rapidly impacts parasite metabolism and is able to form adducts with GSH and other thiols in parasites. In order to test whether this is sufficient to induce oxidative stress in parasites, we measured thiol species following treatment with 30 µM of SFN for 3 h, compared to untreated control. Oxidised glutathione (GSSG) and the GSH precursor, gGlu-Cys, were both significantly elevated in treated cells, while reduced GSH was decreased (Fig 7C). The same trend was seen after treatment with only 15 µM SFN for 3 h (Supp Fig 4B), clearly showing that SFN treatment causes oxidative stress in parasites. We performed pulse activity assays with different concentrations of SFN combined with DHA to look for a potentiating effect. For artemisinin sensitive parasites (Cam3.II^REV^), infected cells were treated with a pulse of SFN (20 µM or 30 µM) and a sub-lethal concentration of DHA (100 nM) for 1 h and parasite survival was assessed in the next cycle. We found significantly decreased survival in parasites treated with the SFN-DHA combinations compared to DHA alone (Fig 7D). For artemisinin resistant parasites (Cam3.II^R539T^), infected cells were pulse treated with SFN (10 µM, 20 µM or 30 µM) and DHA (700 nM) for 3 h. Parasite survival was significantly decreased in parasites treated with the SFN-DHA combinations compared to DHA alone (Fig 7D). It should be noted that SFN alone significantly reduced parasite survival (∼15% reduction for 30 µM), especially in artemisinin sensitive parasites (Fig 7D). Levels of heme species were also assessed, demonstrating that SFN treatment up to 30 µM for 3 h had no impact on hemoglobin, heme or hemozoin levels (Fig 7E and Supp Fig 4C). The rate of drug activation in early ring-stage Cam3.II^R539T^ and Cam3.II^C580Y^ parasites was also measured and revealed no significant difference in DHA activation following either simultaneous treatment with 30 µM of SFN (Fig 7F) or pre-treatment with 15 µM of SFN (Fig 7G). This confirms that SFN does not impact heme-mediated activation of artemisinins, and that decreased levels of reduced thiols are responsible for the potentiating effect on artemisinin potency.

**Figure 7.**
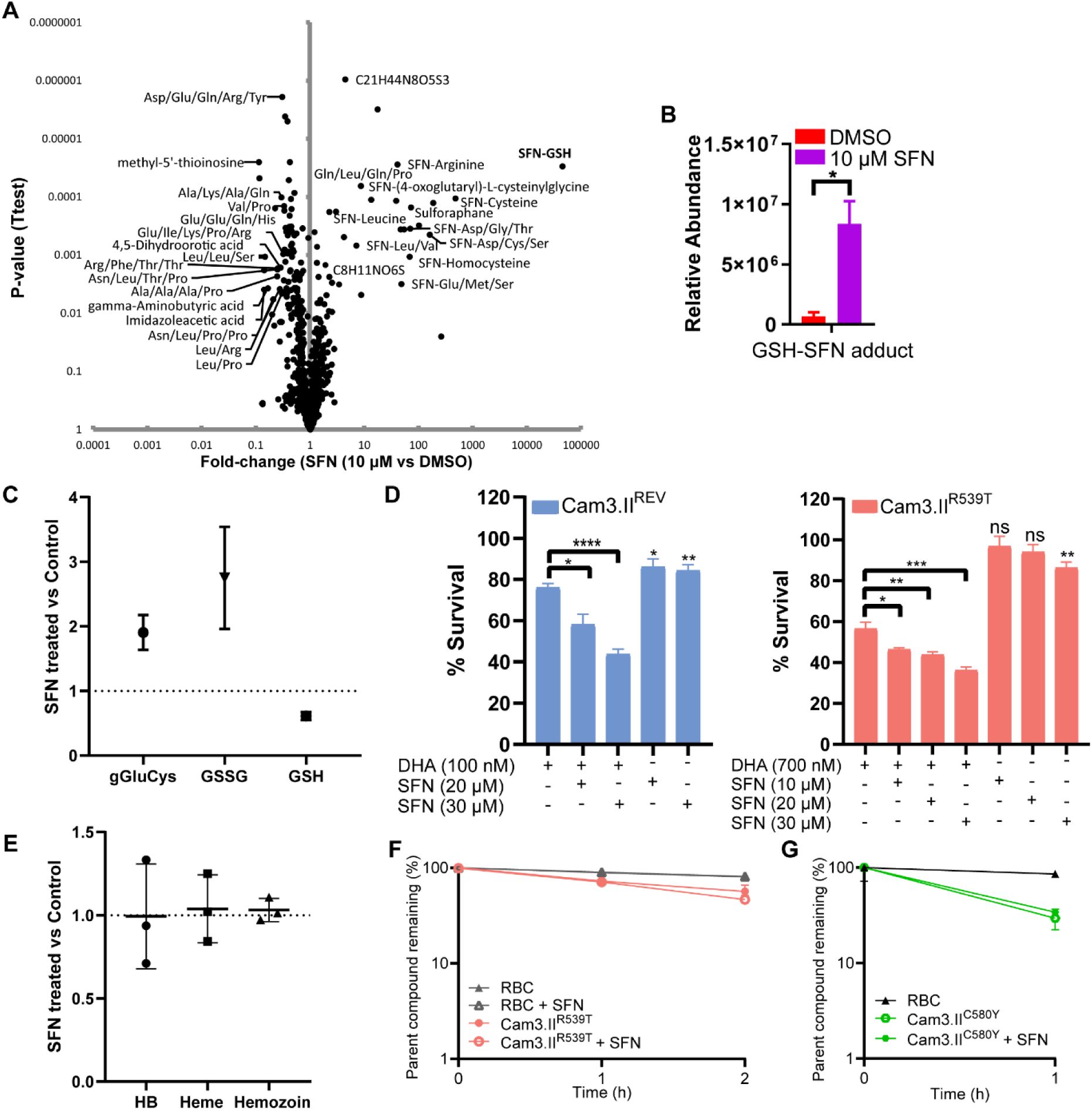
Sulforaphane, a thiol modulating molecule, enhances artemisinin activity. (**A**) Volcano plot showing statistically significant altered metabolites (p-value < 0.01, and fold change (FC) > 3 or < -3) with SFN thiol adducts and putative peptides being the most perturbed metabolites following treatment with 10 µM of SFN for 3 h in Pf3D7 parasites. (**B**) Relative abundance of glutathione (GSH)-sulforaphane (SFN) adduct measured using untargeted metabolomics following treatment with 10 µM of SFN for 3 h. Values represent the means of three independent experiments ± the standard deviation of the mean (SD), *, p-value < 0.05). (**C**) NEM-derivatised γ-glutamyl cysteine (gGlu-Cys), oxidised glutathione (GSSG) and reduced glutathione (GSH) fold change in Pf3D7 wildtype parasites treated with 30 µM SFN for 3 h. Values represent the means of seven independent experiments ± the standard errors of the mean (SEM). (**D**) 3-6 hpi Ring-stage survival assay determination for artemisinin sensitive (Cam3.II^REV^ (blue)) and resistant (Cam3.II^R539T^ (red)) parasites following treatment with either 100 nM of DHA for 1 h (sensitive parasites) or 700 nM of DHA for 3 h (resistant parasites) alone, DHA with increasing doses of SFN (10-30 µM), and SFN alone. Values represent the means of six independent experiments ± the standard errors of the mean (SEM), p-value was calculated using Welch’s t-test (*, p-value < 0.05, **, p-value < 0.01, ***, p-value < 0.001, ****, p-value < 0.0001). DHA (700 nM) degradation profiles in 3-6 hpi ring-stage artemisinin resistant (Cam3.II^R539T^ (red) – 20% parasitemia) (E) (Cam3.II^C580Y^ (green) – 96% parasitemia) parasites and in uninfected red blood cells (gray) following simultaneous treatment with 30 µM SFN. (**E**) Hemoglobin (Hb) and its species (heme and hemozoin) fold-change in 24 hpi trophozoite-stage Pf3D7 wildtype parasites treated with 30 µM SFN for 3 h. Values represent the means of three independent experiments ± the standard deviation of the mean (SD). (**F**) or pre-treatment with 15 µM SFN for 1 h (**G**). Values represent the means of technical replicates from one experiment ± the standard deviation of the mean (SD).

### *In vivo* data shows that sulforaphane can re-sensitise artemisinin resistant parasites

To investigate the potentiating activity of SFN *in vivo*, mice were infected with artesunate sensitive (1804^wt^) or resistant (1804^M488I^) *P. berghei* (ANKA) strains. Once the parasitemia was 2%, mice were injected for 3 consecutive days with 60 mg/kg artesunate, or artesunate combined with 20 mg/kg of SFN and the parasitemia monitored daily. As expected, treatment of 1804^wt^ -infected mice with artesunate was effective, and resulted in only low-level parasite replication after 5-8 days that was not significantly altered in the presence of SFN (Fig 8A). Conversely, the K13-mutant parasite line (1804^M488I^) was less susceptible to artesunate treatment, but the parasite load was significantly reduced in the presence of SFN (Fig 8A) and the mice survived for longer (Fig 8B-C). The artesunate resistance observed with 1804^M488I^ parasites was reduced in the presence of SFN, with parasite loads not significantly different from 1804^wt^ parasites treated with artesunate only (Fig 8A). The dose of SFN administered was titrated down and 5mg/kg SFN was found to be sufficient to significantly prolong the survival of artesunate-treated mice infected with 1804^M488I^ parasites (Fig 8D-E).

**Figure 8.**
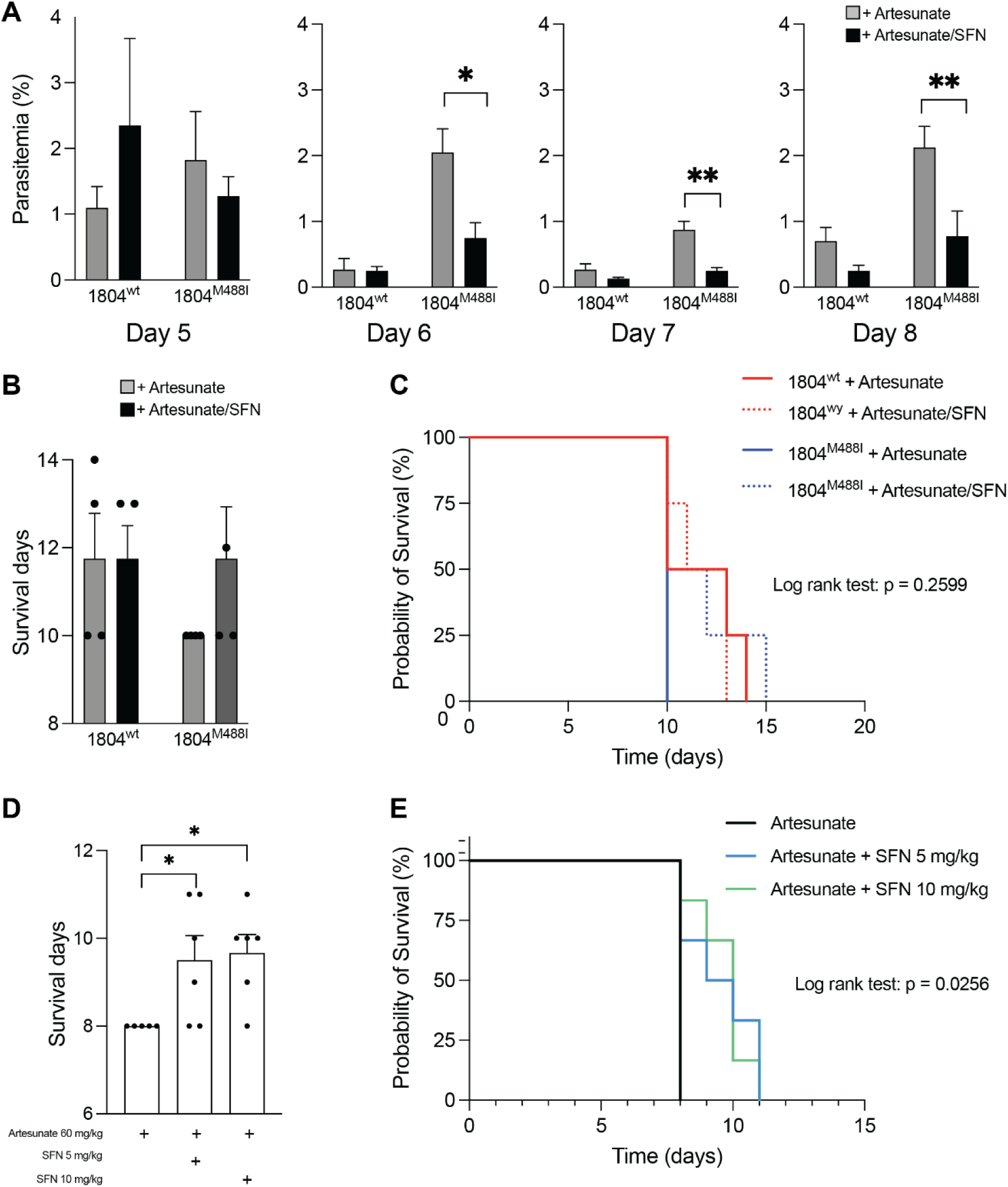
SFN can potentiate artesunate activity *in vivo*. (**A**) Groups of 4 mice were infected with PbANKA sensitive (1804^WT^) and resistant (1804^M488I^) lines on day 0. After 4-6 days, mice were injected on 3 consecutive days with 60mg/kg Artesunate, ± 20mg/kg of SFN and the parasite load measured periodically. Graphs show mean % +/-SEM. (**B**) The average number of days each group of mice survived. Cardiac puncture was performed once the mice showed a parasitemia >15%. No difference was observed for the sensitive lines. (**C**) Kaplan Meier Plot show survival data for each group (n = 4). (**D**) Mice were infected with artesunate resistant strain 1804^M488I.^ Once the infection was established, artesunate was administered on 3 consecutive days with the indicated SFN doses. Both treatment levels prolonged the survival of mice infected with the resistant strain. (**E**) Kaplan Meier Plot show survival data for each group (n = 5-6). Values represent the mean ± standard errors of the mean (SEM), p-value was calculated using Welch’s t-test (*, p-value < 0.05, **, p-value < 0.01, ***, p-value < 0.001, ****, p-value < 0.0001).

## DISCUSSION

Partial resistance to artemisinin has emerged in several countries in Africa and for the prevention of a potential public health disaster, the time for decisive action to confront emerging artemisinin resistance is now. In this study, we have characterised the biological basis of artemisinin resistance and identified a vulnerability in these parasites that can be targeted with available compounds. We showed that PfK13 protein levels are significantly decreased in a range of artemisinin resistant parasites at the early 3-6 hpi ring-stage, which is the stage known to be critical for artemisinin partial resistance. The same trend was observed in trophozoite-stage parasites. This decreased PfK13 abundance correlated with impaired hemoglobin uptake and digestion. In 3-6 hpi ring-stage resistant parasites, we demonstrated lower levels of hemoglobin-derived heme, which is the crucial activator of artemisinin in parasites. Indeed, we have now provided direct evidence that reduced PfK13 levels lead to slower activation of artemisinin in resistant parasites.

Resistant parasites also exhibited higher levels of reduced GSH and its precursor gGlu-Cys, and lower levels of oxidised GSSG, in both 3-6 hpi ring-and trophozoite-stage parasites. We identified this modified parasite redox capacity as a point of vulnerability, which can be targeted to reverse artemisinin resistance. Depleting GSH levels with BSO increased artemisinin sensitivity without affecting heme levels. Increased efflux of GSH in PfMRP1 mutant parasites decreased thiol levels and enhanced artemisinin sensitivity, while increasing reduced GSH levels with NAC decreased artemisinin potency. This clearly supports the impact of thiol levels on artemisinin sensitivity and we then demonstrated that SFN, an available and safe molecule derived from broccoli sprout, can potentiate artemisinin activity both *in vitro* and *in vivo* in artemisinin resistant parasites.

PfK13 mutations leading to decreased PfK13 protein levels, impaired hemoglobin uptake and an elevated antioxidant capacity, are central to artemisinin resistance. Our findings highlight the importance of heme in artemisinin activation and the role of oxidative stress in drug efficacy. We hypothesised that the decreased level of heme from impaired hemoglobin uptake and digestion contributes to maintaining parasites in a reduced state. However, when hemoglobin digestion was inhibited for a short duration (3 h), thiol levels were not affected. On the other hand, when we disrupted hemoglobin uptake using the PfK13 knocksideways line (RAP treatment for 24 h), thiol levels were perturbed. This discrepancy may indicate that the duration of hemoglobin perturbation is key to affecting thiol species, or that it is the uptake of hemoglobin (and other host material) rather than inhibition of digestion that may be key to elevating thiol levels. It could also be that in addition to hemoglobin uptake, PfK13 has an additional role in maintaining parasite redox homeostasis. This is supported by transcriptional profiling of resistant field isolates having changes linked to redox metabolism^20^. Our DIA-MS proteomics analysis of early ring-stage parasites demonstrated that several proteins involved in the parasite respiratory chain, a known source of reactive oxygen species (ROS)^49^, are downregulated in parasites with PfK13 mutations, and since GSH scavenges ROS^23^, less ROS could contribute to the elevated GSH levels in these resistant parasites. Although modest changes in mRNA levels of redox enzymes have previously been reported in PfK13 mutant lines^20^, our proteomics data showed that enzymes involved in GSH synthesis were unaffected by K13 mutations. However, the elevated levels of gGlu-Cys in artemisinin resistant parasites suggest increased activity of the glutathione synthesis pathway, likely driven by a post-translational mechanism.

Based on our findings, PfK13 levels, heme metabolism indicators, peroxide activation rates and/or redox markers could be used to develop field-applicable resistance monitoring tools. Combining multiple resistance markers with genetic data and traditional drug efficacy tests can provide a comprehensive picture of resistance patterns. This is particularly important now that artemisinin partial resistance is emerging in Africa. Resistance to artemisinins in Africa is associated with novel PfK13 mutations and non-PfK13 mutations that have not been studied extensively^50^. It will be important to characterise the role of artemisinin activation rate and redox capacity in the resistance mechanism of these parasites. Techniques like mass spectrometry, while highly accurate, may not be practical for field use, however, these biomarkers could be translated into rapid diagnostic tests for field use. Future work to validate reliable biomarkers and create practical diagnostic assays could enhance our ability to detect and respond to resistance in real-time, ultimately improving malaria control and treatment outcomes, which is absolutely urgent in the current times.

Targeting redox processes and thiol metabolism offers a promising approach to overcoming artemisinin resistance. More specifically in this study, we showed compounds like SFN, which modulate thiol levels, could be used in combination with artemisinin to extend the useful lifespan of artemisinin-based treatments. SFN has antioxidant properties for the host through activation of Nrf2^48^, therefore our molecule of choice would not only kill the parasite, but will boost the host antioxidant capacity. This differs from most other available pro-oxidants, which do not have this host antioxidant capacity. Further, the available human safety and PK data on SFN is highly attractive^51^, and with our *in vitro* and *in vivo* data, this combination could be further developed to determine clinical safety and efficacy in the malaria context. Broccoli sprout extracts are already commercially available and affordable, providing the potential for translation of cost-efficient therapeutics. Many SFN clinical trials are currently ongoing worldwide in various fields including cancer, COVID-19 and vascular disorders of pregnancy^52,53^. However, consideration needs to be taken around SFN product quality and availability as an approved medicine rather than a nutritional supplement in malaria endemic regions. In the future, this limitation is likely to be addressed by the availability of stabilised synthetic pharmaceutical grade SFN (e.g. Sulforadex®), which has been developed and is being tested in clinical trials for a range of different cancers. Using SFN in combination with artemisinin presents a novel and promising strategy to tackle artemisinin resistance. By leveraging SFN’s ability to induce oxidative stress and deplete thiol levels in parasites, this approach can enhance the efficacy of artemisinin and potentially restore its effectiveness against resistant strains. Further research and clinical studies will be required to translate these findings into practical treatment regimens that can be widely adopted in the fight against malaria.

## Acknowledgments

We would like to acknowledge the traditional custodians of the lands this project was conducted on: the Wurundjeri people of the Kulin nation. Metabolomics and Proteomics analysis was conducted with assistance from Dovile Anderson and the Monash Proteomics and Metabolomics Platform. We thank the Australian Red Cross LifeBlood Service for provision of RBCs. We thank Andy Waters for providing the Pb ANKA parasite lines, Tobi Spiellman for providing the PfK13 knocksideways line, David Fidock for providing the Cam3.II lines, Leann Tilley for providing the PL2 and PL7 lines, and the Malaria Research and Reference Reagent Resource Centre (MR4) for providing the MRA lines. DJC acknowledges funding from an NHMRC Career Development Fellowship (II) #APP1148700, ARC Future Fellowship (#FT220100564) and NHMRC Synergy Grant (#APP1185354). NC acknowledges funding from Deakin University IMPACT SEED Grant.

The manuscript was written through contributions of all authors. All authors have given approval to the final version of the manuscript. The authors declare no conflict of interest. Proteomics data are available via ProteomeXchange with identifier PXD068108. Metabolomics data is available at the NIH Common Fund’s National Metabolomics Data Repository (NMDR) Metabolomics Workbench with Project ID ST004194.

