## Supplementary Figures and Table for "PfK13-associated artemisinin resistance slows drug activation and enhances antioxidant defence, which can be overcome with sulforaphane"

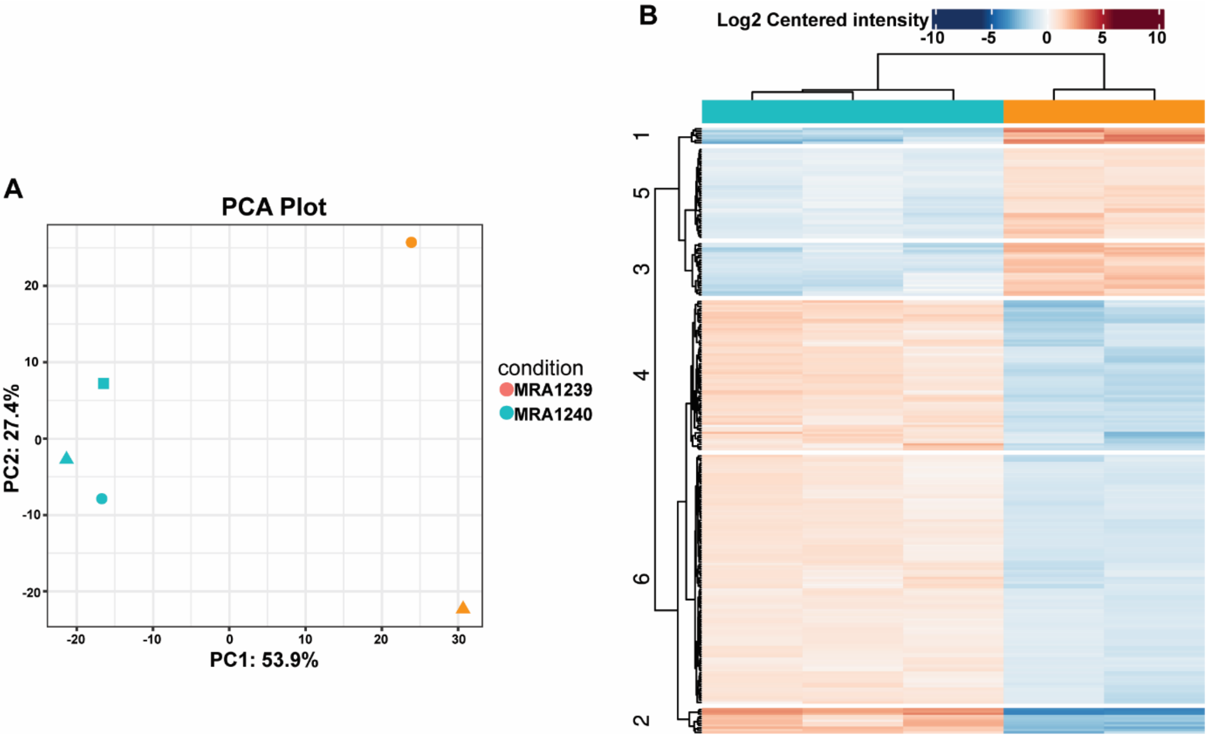


**Supplementary Figure S1. Proteomics analysis of artemisinin resistant (MRA1240) and sensitive (MRA1239) parasites. (A**) Principal component analysis of all the 2407 proteins quantified across MRA1240 (artemisinin resistant and 3 samples) and MRA1239 (artemisinin sensitive and 2 samples). (**B**) Hierarchical clustering analysis of significantly dysregulated proteins across the conditions (light blue – MRA1240, orange – MRA1239) (log2 fold-change of 0.5 and Bonferroni corrected p-value of less than 0.05).

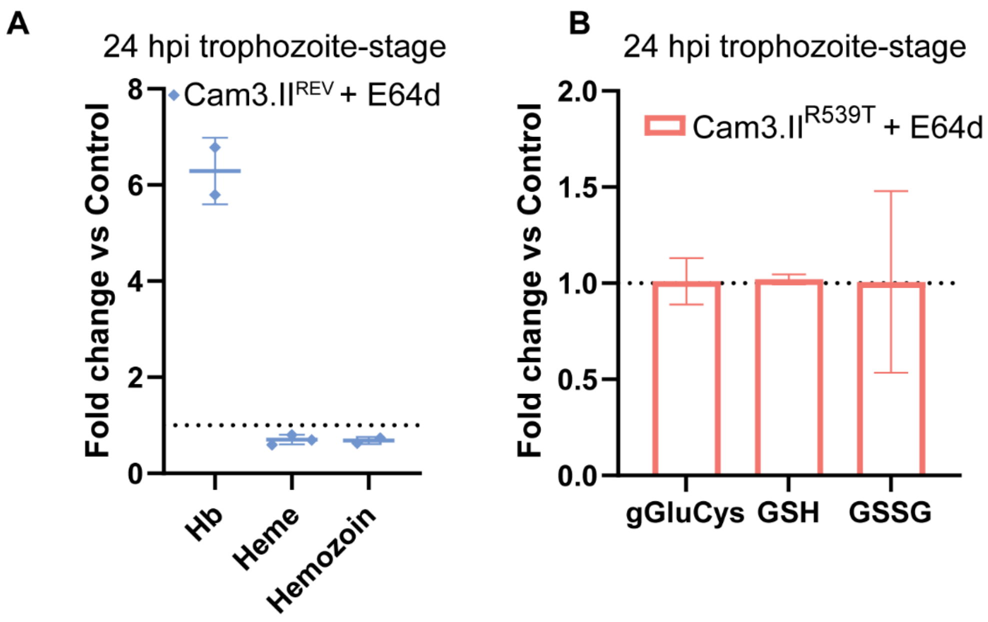


**Supplementary Figure S2. Modulating hemoglobin catabolism does not affect thiol levels.** (**A**) Hemoglobin (Hb) and its species (heme and hemozoin) fold-change in 24 hpi trophozoite-stage artemisinin-sensitive (Cam3.II$^{REV}$) parasites treated with 10 µM E64d for 3 h. Values represent the means of two-three independent experiments ± the standard deviation of the mean (SD). (**B**) NEM-derivatised γ-glutamyl cysteine (gGlu-Cys), reduced glutathione (GSH) and oxidised glutathione (GSSG) fold change in artemisinin resistant parasites treated with 10 µM E64d for 3 h. Values represent the means of three independent experiments ± the standard errors of the mean (SEM).

**A** **Chromosome 1: amplified genes**

| Gene ID | Gene description |
|---|---|
| PfDd2_010015700 | multidrug resistance-associated protein 1 |
| PfDd2_010015800:rRNA | rRNA |
| PfDd2_010015900:rRNA | rRNA |
| PfDd2_010016000:rRNA | rRNA |
| PfDd2_010016300 | Plasmodium exported protein (hyp11), unknown function |
| PfDd2_010016400 | Plasmodium exported protein, unknown function |
| PfDd2_010016500 | glutamic acid-rich protein |
| PfDd2_010016600 | surface-associated interspersed protein 1.1 (SURFIN 1.1) |
| PfDd2_010016700 | Plasmodium exported protein, unknown function |
| PfDd2_010016800 | Plasmodium exported protein (hyp1), unknown function |
| PfDd2_010016900 | Plasmodium exported protein, unknown function |
| PfDd2_010017000 | surface-associated interspersed protein 1.2 (SURFIN 1.2) |
| PfDd2_010017100 | heat shock protein 40, type II |
| PfDd2_010017200 | DBL containing protein, unknown function |
| PfDd2_010017300 | Plasmodium exported protein (hyp8), unknown function |
| PfDd2_010017400 | exported protein family 1 |
| PfDd2_010017500 | Pfmc-2TM Maurer's cleft two transmembrane protein |
| PfDd2_010017600 | exported protein family 3 |
| PfDd2_010017700 | exported protein family 4, pseudogene |
| PfDd2_010017900 | erythrocyte membrane protein 1 (PfEMP1), exon 2, pseudogene |
| PfDd2_010018000 | Plasmodium exported protein (hyp10), unknown function |
| PfDd2_010018100 | stevor, pseudogene |
| PfDd2_010018200 | rRifin |
| PfDd2_010018300 | Plasmodium exported protein (hyp7), unknown function |
| PfDd2_010018400 | Plasmodium exported protein, unknown function, pseudogene |
| PfDd2_010018500 | surface-associated interspersed protein 1.3 (SURFIN 1.3) |
| PfDd2_010018750 | Plasmodium exported protein, unknown function, pseudogene |
| PfDd2_010018800 | Rifin |
| PfDd2_010018900 | Rifin |
| PfDd2_010019000 | Rifin |
| PfDd2_010019100 | Rifin |
| PfDd2_010019200 | Rifin |
| PfDd2_010019300 | Rifin |
| PfDd2_010019400 | erythrocyte membrane protein 1, PfEMP1 |
| PfDd2_010019500 | erythrocyte membrane protein 1 (PfEMP1), exon 2 |
| PfDd2_010019550 | erythrocyte membrane protein 1 (PfEMP1), exon 2 |

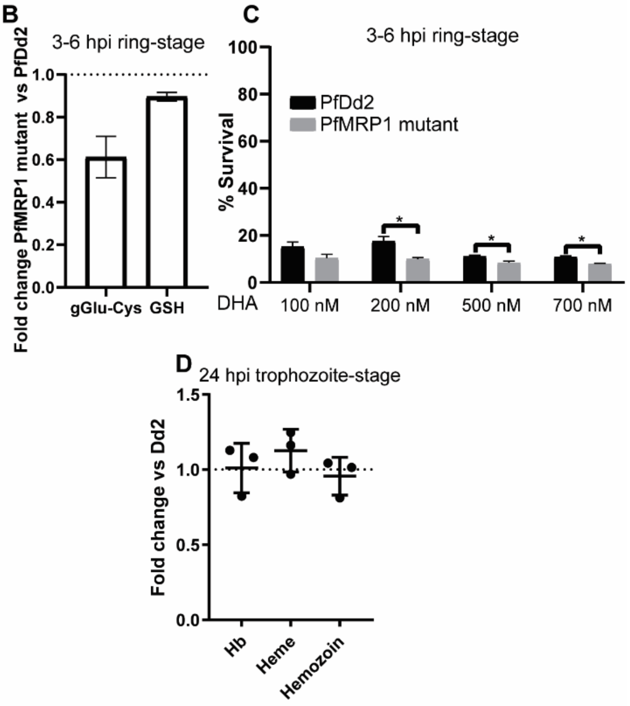


**Supplementary Figure S3.** ***Plasmodium falciparum*** **multidrug resistance protein 1 (PfMRP1) line has decreased glutathione levels, which affects artemisinin sensitivity.** **(A)** List of amplified genes from chromosome 1 in PfMRP1 mutant, with PlasmoDB identifier and gene description listed. **(B)** NEM-derivatised gGlu-Cys and GSH fold change in PfMRP1 mutant compared to PfDd2 control. Values represent the means of three independent experiments ± the standard errors of the mean (SEM). **(C)** 3-6 hpi ring-stage survival assay determination for PfDd2 wildtype and PfMRP1 mutant parasites following treatment with different doses of DHA for 1 h. Values represent the means of two independent experiments ± the standard deviation of the mean (SD), p-value was calculated using Welch's t-test (*, p-value < 0.05). **(D)** Hb and its products (heme and hemozoin) relative abundance (absorbance measured) in 24 hpi trophozoite-stage PfDd2 and PfMRP1 mutant parasites. Values represent the means of three technical replicates ± the standard deviation of the mean (SD).

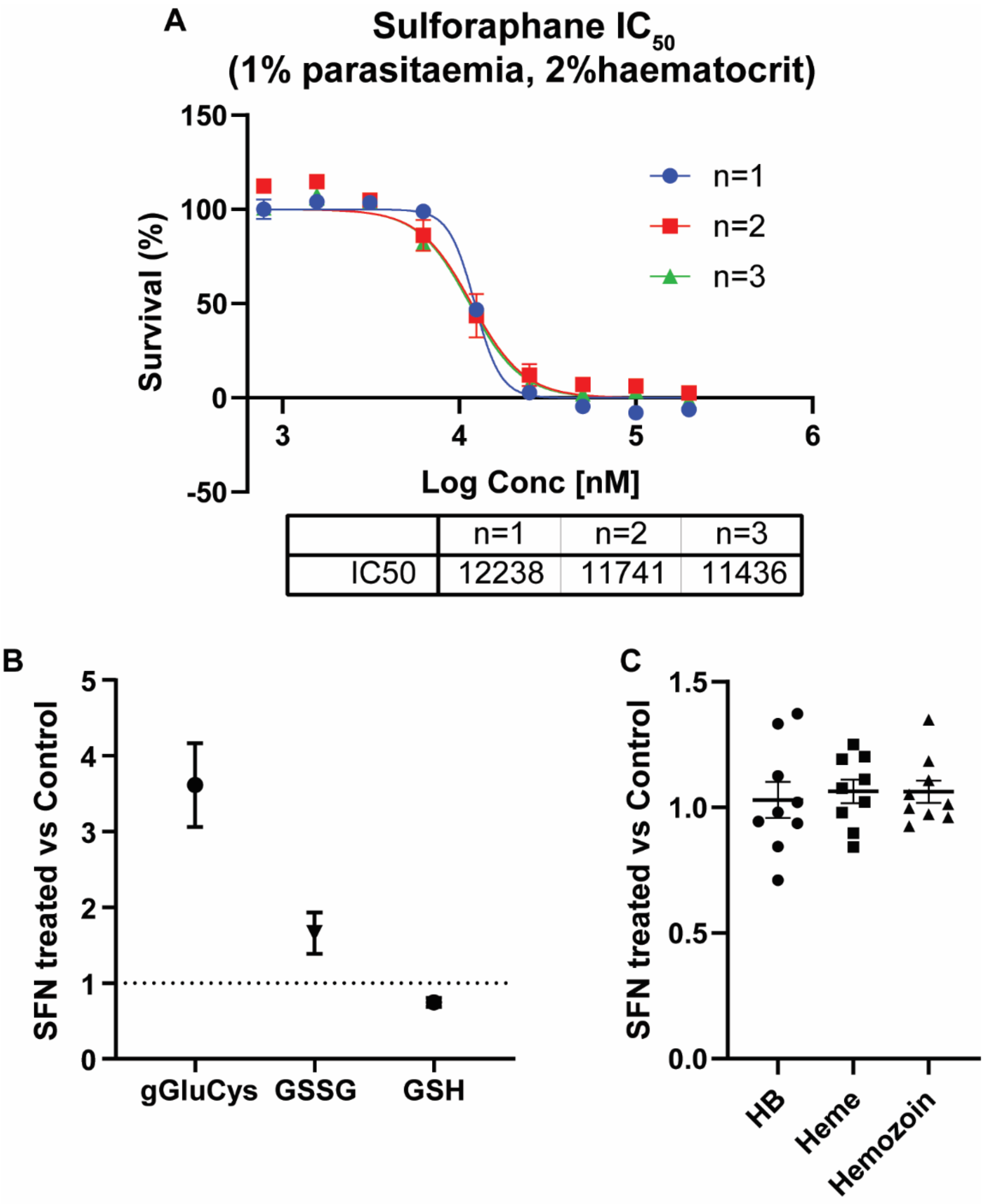


**Supplementary Figure S4. Sulforaphane affects parasite thiol species but not heme levels.** (**A**) $IC_{50}$ curves (from three biological replicates) of parasites treated with SFN were determined using the SYBR Green assay. Each data point represents the mean of two technical replicates ± SD. The $IC_{50}$ of SFN was 11.8 ± 0.4 μM. (**B**) NEM-derivatised γ-glutamyl cysteine (gGlu-Cys), oxidised glutathione (GSSG) and reduced glutathione (GSH)

fold change in Pf3D7 wildtype parasites treated with 15 µM SFN for 3 h. Values represent the means of three independent experiments ± the standard errors of the mean (SEM). (**C**) Hemoglobin (Hb) and its products (heme and hemozoin) fold-change in 24 hpi trophozoite-stage Pf3D7 wildtype parasites treated with 15 µM SFN for 3 h. Values represent the means of ten independent experiments ± the standard errors of the mean (SEM).

**Supplementary Materials:**

**Supplementary Data Sheet 1 -** Protein intensities for DIA experiments - Artemisinin resistant (Cam3.II$^{R539T}$ and Cam3.II$^{C580Y}$) Vs sensitive (Cam3.II$^{REV}$).
**Supplementary Data Sheet 2 -** Protein intensities for DIA experiments - Artemisinin resistant (MRA1240) Vs sensitive (MRA1239).
**Supplementary Data Sheet 3 -** IDEOM metabolomics output for Sulforaphane treatment of *P. falciparum* (3D7 wildtype strain).

**Supplementary Table 1.** List of significantly dysregulated parasite proteins (Log2 Fold-change of 0.5 and P-value < 0.05) in 6 hpi ring-stage artemisinin resistant compared to artemisinin sensitive parasites. In addition, their importance in parasite growth and proteomics expression profile across the asexual stage, whether interaction with PfK13 is observed using BioID and pulldown experiments, differentially regulated in two published ring-stage proteomics (not 6 hpi) and a 24 hr trophozoite-stage proteomics study (using data-independent acquisition).

| **Genes ID/name** | **Protein Descriptions** | **Importance for growth** | **Log$_2$ Fold change Cam3.II$^{R539T}$ vs Cam3.II$^{REV}$** | **Log2 Fold change Cam3.II$^{C580Y}$ vs Cam3.II$^{REV}$** | **p-value Cam3.II$^{R539T}$ vs Cam3.II$^{REV}$** | **p-value Cam3.II$^{C580Y}$ vs Cam3.II$^{REV}$** | **Proteomics expression across the red blood cell stage (6 h Rings - 24 h Trophozoites and 38 h - Schizonts) in DIA Siddiqui *et al.* 2022[30]** | | | **Birnbaum *et al.* 2020[8] PfK13 Bio-ID[8]** | **Gnadig *et al.* 2020[15] PfK13 pulldown[15]** | **Differentially regulated in Gnadig *et al.* 2020[15] ring-stage proteomics** | **Differentially regulated in DDA Siddiqui *et a.* 2017[9] ring-stage proteomics (Cam3.II$^{R539T}$ vs Cam3.II$^{REV}$)** | **Differentially regulated in DIA Siddiqui *et al.* 2022[30] trophozoite-stage proteomics Cam3.II$^{R539T}$ vs Cam3.II$^{REV}$)** |
|---|---|---|---|---|---|---|---|---|---|---|---|---|---|---|
| | | | | | | | Rings | Trophozoite | Schizonts | | | | | |
| PF3D7_0310500 | ATP-dependent RNA helicase DHX57, putative | not essential | -2.14 | -1.91 | 0.0031 | 0.0051 | 5.149 | 5.670 | 5.625 | | | | | |
| PF3D7_0320800 | ATP-dependent RNA helicase DDX6 | not essential | -2.00 | -1.40 | 0.0126 | 0.0179 | 5.637 | 5.917 | 5.884 | Enriched | | | | Decreased |
| PF3D7_0401800 | Plasmodium exported protein (PHISTb), unknown function | not essential | -4.62 | -4.59 | 0.0293 | 0.0290 | 5.904 | 6.512 | 6.596 | Enriched | | | Decreased (1 of 2 biological reps) | Decreased |
| PF3D7_0407200 | peptidyl-tRNA hydrolase 2, putative | essential | -2.26 | -3.75 | 0.0070 | 0.0003 | 5.031 | 4.895 | 4.775 | | | | Decreased* | Increased |
| PF3D7_0422300 | alpha tubulin 2 | not essential | -1.87 | -2.29 | 0.0011 | 0.0079 | NA | | | | | | | |
| PF3D7_0603500 | cation/H+ antiporter | not essential | -4.06 | -4.01 | 0.0246 | 0.0251 | 4.771 | 4.872 | 5.168 | | | | | |
| PF3D7_0708100 | DNA-directed RNA polymerases I, II, and III subunit RPABC5, putative | essential | -2.68 | -1.47 | 0.0343 | 0.0449 | 5.412 | 5.518 | 5.424 | | | | | |
| PF3D7_0708900 | protein SCO1, putative | essential | -3.47 | -3.39 | 0.0113 | 0.0123 | 4.311 | 4.476 | 4.259 | | | | | |
| PF3D7_0820700 | 2-oxoglutarate dehydrogenase E1 component | not essential | -2.93 | -2.88 | 0.0072 | 0.0072 | 5.116 | 5.217 | 5.226 | | Enriched | | Decreased* | |
| PF3D7_0823300 | histone acetyltransferase GCN5 | essential | -1.98 | -2.06 | 0.0002 | 0.0008 | 5.352 | 5.551 | 5.531 | | | | | |
| PF3D7_0929900 | conserved Plasmodium protein, unknown function | essential | -3.46 | -2.05 | 0.0186 | 0.0236 | 3.877 | 4.536 | 4.436 | | | | | |
| PF3D7_0936800 | Plasmodium exported protein (PHISTc), unknown function | not essential | -1.53 | -2.18 | 0.0399 | 0.0430 | 5.959 | 6.106 | 6.120 | Enriched | | | | |

| | | | | | | | | | | | | | | |
|---|---|---|---|---|---|---|---|---|---|---|---|---|---|---|
| PF3D7_1021900 | PHAX domain-containing protein, putative | essential | -3.21 | -3.10 | 0.0168 | 0.0238 | 5.601 | 6.194 | 6.301 | | | | | |
| PF3D7_1024300 | ATP synthase-associated protein, putative | essential | -1.62 | -1.83 | 0.0427 | 0.0333 | NA | | | | | | | |
| PF3D7_1129900 | major facilitator superfamily-related transporter, putative | not essential | -1.80 | -2.74 | 0.0162 | 0.0250 | 5.367 | 5.703 | 5.571 | | | | Decreased* | |
| PF3D7_1212800 | succinate dehydrogenase [ubiquinone] iron-sulfur subunit, mitochondrial | not essential | -2.37 | -2.30 | 0.0097 | 0.0062 | 4.692 | 4.718 | 5.012 | | | | | |
| PF3D7_1219600 | phospholipid-transporting ATPase 2 | essential | -1.72 | -1.76 | 0.0101 | 0.0121 | 5.258 | 5.185 | 5.230 | Enriched | | | | |
| PF3D7_1237200 | conserved Plasmodium protein, unknown function | not essential | -1.79 | -2.00 | 0.0008 | 0.0017 | 4.738 | 5.329 | 5.193 | | | | | |
| PF3D7_1248700 | conserved protein, unknown function | essential | -2.13 | -1.78 | 0.0044 | 0.0112 | 5.453 | 5.654 | 5.816 | | | | | |
| **PF3D7_1343700** | **kelch protein K13** | **essential** | **-1.44** | **-1.32** | **0.0277** | **0.0281** | **6.588** | **6.312** | **6.405** | **Enriched** | **Enriched** | **Significantly decreased only in Cam3.II$^{R539T}$ compare to control** | **Decreased** | **Decreased** |
| PF3D7_1361900 | proliferating cell nuclear antigen 1 | essential | 1.87 | 1.36 | 0.0474 | 0.0292 | 6.918 | 6.983 | 7.0848 | Enriched | | | | |
| PF3D7_1403900 | serine/threonine protein phosphatase CPPED1, putative | essential | 2.18 | 1.18 | 0.0161 | 0.0269 | 5.395 | 5.540 | 5.461 | | | Significantly increased in resistant parasites | | |
| PF3D7_1439400 | cytochrome b-c1 complex subunit Rieske, putative | not essential | -1.45 | -1.75 | 0.0431 | 0.0455 | 4.962 | 5.130 | 5.141 | | Enriched | | Decreased* | |
| PF3D7_1441400 | FACT complex subunit SSRP1, putative | essential | -1.67 | -1.21 | 0.0244 | 0.0458 | 6.390 | 6.412 | 6.432 | Enriched | | | Decreased (2 of 3 biological reps) | |
| PF3D7_1454400 | aminopeptidase P | essential | 2.18 | 1.50 | 0.0122 | 0.0233 | 6.691 | 7.090 | 6.956 | Enriched | | Significantly increased in resistant parasites | Increased (1 of 3 biological reps) | |
| PF3D7_1466800 | NOC3 domain-containing protein, putative | essential | -2.61 | -1.82 | 0.0433 | 0.0414 | 4.717 | 5.082 | 4.688 | Enriched | | | Decreased* | |
| PF3D7_1478000 | Plasmodium exported protein (PHISTa), unknown function | essential | -1.76 | -2.45 | 0.0338 | 0.0334 | NA | | | | | | | |

*detected in one experiment
